# Whole-genomes illuminate the drivers of gene tree discordance and the tempo of tinamou diversification (Aves: Tinamidae)

**DOI:** 10.1101/2024.01.22.576737

**Authors:** Lukas J. Musher, Therese A. Catanach, Thomas Valqui, Robb T. Brumfield, Alexandre Aleixo, Kevin P. Johnson, Jason D. Weckstein

## Abstract

As an old group that has diversified in South America over millions of years, the tinamous (Palaeognathae: Tinamidae) are of high interest for understanding the evolution of birds and the assembly of the Neotropical biota. However, there are currently no complete species-level phylogenies of this group. Most prior work has been based on either morphological data or a small number of molecular markers, each of which has limited capability for reconstructing the tinamou phylogeny. Therefore, the interrelationships of most tinamou species are uncertain. We analyzed 80 whole-genomes from a mix of historical study skins and frozen tissues, including all 46 recognized species of tinamous to (1) reconstruct their interrelationships, (2) estimate the timeframe of tinamou evolution, and (3) examine for the effects of incomplete lineage sorting (ILS) and ancestral introgression on genome evolution. We compared results for coding (BUSCO) and ultraconserved element (UCE) loci, as well as sex-linked and autosomal markers, and used fossil-calibrated tip-dating to estimate divergence times. Tinamous diverged from their sister-group, the extinct Moas, 50-60 mya, and their crown divergence occurred roughly 30-40 mya, followed by constant diversification rates until the present. Phylogenetic reconstructions were largely robust across methods and datasets. Only one clade in the genus *Crypturellus* displayed substantial species-tree discordance across the different data sets. To investigate the impacts of introgression on this discordance, we quantified introgression for 100kb non-overlapping windows across the genome, and identified pervasive genome-wide introgression. The distribution of this introgression across the genome was dependent on the assumed phylogeny applied to the f-branch model. When assuming one of these topologies in the f-branch model, patterns of introgression matched theoretical predictions about genome architecture. Overall, we present the most complete phylogeny for tinamous to date, identify an unrecognized species, and provide a case study for species-level phylogenomic analysis using whole-genomes.

## Introduction

Neotropical organisms have been at the center of discussions about the drivers of the diversification and assembly of tropical biotas (Hughes and Eastwood 2006; Antonelli et al. 2018; Albert et al. 2020; Capurucho et al. 2023). However, most groups lack complete species-level phylogenies, which are crucial for testing questions in these fields. To this end, the tinamous (Palaeognathae: Tinamidae) are of key interest. First, as the only flighted group of palaeognathous birds, their phylogenetic placement among this group is an enigma (Cracraft 1974; Harshman et al. 2008; Cloutier et al. 2019). Second, they are relatively old, having been isolated in Neotropical regions for at least 25 million years (Claramunt and Cracraft 2015; Prum et al. 2015; Yonezawa et al. 2017). However, there has been limited work on tinamou phylogenomics to date. There are no complete species-level tinamou phylogenies, and prior phylogenetic studies utilized either phenotypic data with few-to-no genetic markers (Bertelli and Porzecanski; Bertelli et al. 2002; Valqui 2009; Bertelli 2017; Almeida et al. 2022), or phylogenomic data with minimal taxonomic sampling (Cloutier et al. 2019). Thus, the species-level relationships of the tinamous remain uncertain, hampering efforts to understand the diversification of this enigmatic group.

Unfortunately, obtaining even a near-complete and robust phylogeny for tinamous represents a considerable challenge. Although there has been a pronounced effort to collect modern voucher-linked fresh tissues of Neotropical birds (Moncrieff et al. 2019, 2021; Del-Rio et al. 2021), by comparison to other groups, tinamous are poorly represented in modern tissue collections. Thus, a mix of historical samples with degraded DNA and intraspecific sampling is needed to better document tinamou diversity and phylogenetic interrelationships (McCormack et al. 2016). This is important because some studies have suggested that currently recognized genera are non-monophyletic, warranting further study to reevaluate their taxonomic treatment (Bertelli 2017; Almeida et al. 2022). Moreover, and in line with other Neotropical bird groups, current taxonomic treatments may also misrepresent true species-level diversity (Krabbe et al. 2020; Freeman and Pennell 2021; Musher et al. 2023). Deep divergences and non-reciprocal monophyly (i.e., paraphyly/polyphyly) of intraspecific populations suggest that some widespread species likely include multiple species-level taxa by some criteria (Valqui 2009).

The lack of species-level phylogenies for tinamous also hampers efforts to understand the timeframe of diversification of this group. Past studies have concluded that the crown divergence within tinamous resulted in two major clades (subfamilies), the forest dwelling Tinaminae and open country Nothurinae. However, current estimates for this node vary from roughly 20 mya to as much as ∼50 mya (Mitchell et al. 2014; Prum et al. 2015; Bertelli 2017; Berv and Field 2018; Almeida et al. 2022; Stiller et al. 2024). Moreover, the origins of all extant tinamou genera lack sufficient divergence time estimates. Despite these limitations, there exist multiple crown fossils from both major clades, making this group a good candidate for tip-based dating approaches (Stadler 2010; Didier et al. 2012). An advantage of tip-dating over standard node-calibration is that it does not require the precise phylogenetic placement of fossils into the tree. This is important because the phylogenetic positions of multiple tinamou fossils, such as *Crypturellus reai* Chandler 2012 and *Nothura parvula* Brodkorb 1963, have been equivocal (Bertelli et al. 2014; Almeida et al. 2022). Thus, tip-dating methods allow for the inclusion of these fossils for calibration despite uncertainty about their phylogenetic affinities.

When combined with fossil data, high-throughput sequencing of historical and modern material offers a path forward for obtaining a complete and robust dated phylogeny. Yet even these large datasets, although highly informative, present unique problems (Reddy et al. 2017; Zhao et al. 2023). For example, analyses that utilize alternative datasets or methodological frameworks to answer the same phylogenetic questions have often yielded conflicting, well-supported results (Prum et al. 2015; Simion et al. 2017; Franz et al. 2019; Schultz et al. 2023; Stiller et al. 2024). Some uncertainty arises in the presence of rapid diversification that increases the rate of incomplete lineage sorting (ILS) and likelihood of ancestral introgression, two major biological sources of discordance among data partitions in phylogenetic datasets. These processes increase discordance of gene histories, posing challenges for phylogenetic reconstruction but also offering insights into complex evolutionary processes (Burbrink and Gehara 2018; Rivas-González et al. 2023). An additional challenge for reconstructing rapidly diversifying clades is the paucity of polymorphic nucleotide sites needed for reconstructing short internodes (Leaché et al. 2015; Mclean et al. 2019). Briefly, rapid divergence events can be difficult to reconstruct because the short timeframe also means there is limited phylogenetic signal for those divergence events within individual gene alignments. Interestingly, the use of sex-linked markers may help alleviate these issues as they are known to (1) evolve faster, thus accumulating more substitutions that may be useful for reconstructing recalcitrant nodes, (2) have smaller effective population sizes, thereby reducing rates of gene tree discordance (ILS), and (3) experience lower rates of recombination and introgression (Charlesworth et al. 1987; Meisel and Connallon 2013; Fontaine et al. 2015; Li et al. 2019; Musher et al. 2024). Thus, if genealogical signal across much of the genome is associated with introgression due to ancestral hybridization, but not the bifurcating history of species divergence (i.e., the species tree), sex-chromosomes may be enriched with the signal of the species tree topology (Fontaine et al. 2015; Li et al. 2019; Burbrink et al. 2025).

Here, we sequenced whole genomes for all described tinamou species and many subspecies using a combination of frozen tissues and antique DNA sampled from historical museum skins. Specifically, we reconstruct the evolutionary history in this group while examining and testing for the impacts ILS and introgression on phylogenetic inference. Our objectives are to (1) reconstruct the species-level interrelationships of tinamous, (2) estimate the timeframe of tinamou diversification, and (3) test for the effects of ILS and introgression on genome evolution, including differences between sex-linked and autosomal markers.

## Materials and methods

### Sampling

There are 46 currently recognized species of tinamous classified into nine genera (Clements et al. 2023). Twenty-two of these species are monotypic and the remaining species contain two or more subspecies, totaling 175 named taxa (monotypic species plus subspecies) recognized in Tinamidae. We sampled 55 frozen tissues, 14 toe pads from museum study skins, and downloaded 11 whole-genome assemblies from NCBI Assembly Archive, spanning a total of 71 named taxa and all 46 recognized tinamou species (Table S1).

Tinamous belong to the infraclass, Palaeognathae which includes many extant flightless ratites such as ostriches (Struthionidae) and rheas (Rheidae) along with extinct forms such as Moas (Dinornithiformes). Recent work has corroborated that moas are the sister group to tinamous and that rheas belong to a clade that is sister to tinamous plus moas (Phillips et al. 2010; Baker et al. 2014; Cloutier et al. 2019). Thus, as outgroup taxa for tree rooting, we also downloaded whole genome assemblies for *Rhea pennata* (greater rhea) and *Anomalopteryx didiformis* (little bush moa) from the NCBI Assembly Archive. Table S1 lists the 82 (79 Tinamous + two outgroup taxa) samples included in this study.

### Whole-genome assembly and ortholog identification

The details of whole-genome sequencing and assembly can be found in the Supplementary Materials document. After sequencing, we cleaned and mapped raw reads for each sample (Table S1) to a closely related scaffold-level assembly from NCBI. Decisions about which genome to use for reference mapping were made based on a previously published tinamou phylogeny (Almeida et al. 2022). We used (1) fastp (Chen et al. 2018) to clean raw reads, trim adapters and remove low quality reads, (2) Burrows-Wheeler Aligner (bwa) v0.7.17 (Li and Durbin 2009) to map the cleaned reads to reference genomes, (3) samtools v1.6 to convert the resulting sam-files into sorted bam-files (Li et al. 2009), and (4) Analysis of Next Generation Sequencing Data (ANGSD) v1.2.13 (Korneliussen et al. 2014) to convert bam-files into fasta format genomes. To obtain pseudo-chromosome assemblies for each sample, we then used ragtag v2.1.0 (Alonge et al. 2022) to scaffold each assembly against the chromosome-level *Rhea pennata* genome (NCBI refseq assembly: GCF_028389875.1) using default settings.

We identified and extracted two types of orthologous markers from the whole-genomes. First, we utilized nucleotide sequences of protein-coding genes harvested using Benchmarking of Universal Single Copy Orthologs (BUSCO) v5.3.0 (Simão et al. 2015). We used custom scripts to append orthologous genes from all samples into the same text file before aligning. Second, we harvested ultraconserved elements (UCEs) from each genome assembly, including the entire conserved UCE core plus 1,000 bp of flanking sequence on both 5’ and 3’ ends. UCEs are conserved, putatively single-copy, sequences in the genome that are flanked by sites that become more variable moving away from the conserved core (Faircloth et al. 2012; McCormack et al. 2012). We followed the pipeline outlined in Phyluce v1.7.1 (Faircloth 2016) for harvesting loci from whole genomes using the Tetrapods-UCE-5kv1 probeset of 5,060 UCE’s (see Supplementary Materials). To examine differences among sex-linked and autosomal markers, we also created two datasets of UCE’s (see Supplementary Materials), one dataset of sequences mapping to the Z-chromosome (an avian sex chromosome with idealized N_e_ of 0.75 autosomal N_e_) and one that mapped to autosomes. Some analyses suggested that UCE’s were more reliable markers than BUSCO’s, which had a potential impact of systematic error (see Supplemental Materials); for this reason, we only separated the UCE dataset, and not the BUSCO’s, into sex-linked and autosomal datasets. All datasets were aligned using MAFFT v7.453 (Katoh and Standley 2013).

To evaluate impacts of missing data, we filtered UCE and BUSCO alignments twice, once to keep only those loci with a minimum of 75% of samples represented in each alignment (75% complete datasets) and once to keep only those with complete sampling (100% complete datasets). However, for autosomal and Z-linked datasets, we kept only 75% complete alignments to maximize alignment inclusion in these two smaller datasets. Finally, each alignment was trimmed using TrimAL v1.4 (Capella-Gutierrez et al. 2009) with a gap threshold of 97.5% to trim out potential BUSCO mis-annotations unique to a small number of samples (see Supplementary Materials). This resulted in six primary datasets of sequence alignments used in our analyses for the entire Tinamou dataset (four additional datasets are discussed in the Supplemental Materials document): (1) 75% complete BUSCO, (2) 100% complete BUSCO, (3) 75% complete UCE, (4) 100% complete UCE, (5) 75% complete autosomal UCE, and (6) 75% complete Z-linked UCE.

### Phylogenomic analyses

For each of the six datasets, we employed both a concatenation approach using IQ-TREE v2.0.3 (Minh et al. 2020) and a multi-species coalescent (MSC) approach using ASTRAL-III v5.7.7 (Mirarab et al. 2014; Zhang et al. 2018). For our concatenation approach, we used IQ-TREE to concatenate all alignments and used the edge-linked proportional partition model to partition by locus (Chernomor et al. 2016). In IQ-TREE, we implemented ModelFinder (Kalyaanamoorthy et al. 2017) to choose the best substitution model from the standard model set for each partition from all possible models. We then used IQ-TREE to reconstruct the phylogeny and perform 1,000 ultrafast bootstrap replicates (Hoang et al. 2018).

Next, we estimated phylogenies for each alignment (gene trees) within each of the six datasets, again using IQ-TREE with ModelFinder to identify the best-fit substitution model, and again performed 1,000 ultrafast bootstraps on each gene tree. We then collapsed all low-support (bootstrap<75) nodes within each gene tree (Simmons and Gatesy 2021). Using these gene trees, we implemented a gene tree summarization quartet-based MSC approach in ASTRAL-III. This approach assumes that all gene tree discordance is due to ILS without significant ancestral introgression, recombination, or gene tree estimation error.

In addition to estimating node bootstrap support in IQ-TREE and local posterior probability in ASTRAL-III, we assessed node support using gene concordance factors (gCF’s) for all species trees using IQ-TREE. gCF’s are the percentage of gene trees that contain a given node in a species tree.

### Estimating the tempo of tinamou diversification

To estimate the timeframe of tinamou diversification, we used a fossilized birth-death model with six crown Tinamidae tip-fossil calibrations using an optimized relaxed clock in Beast v2.7.7 with the Sampled Ancestors and Optimized Relaxed Clock packages (Drummond et al. 2006; Gavryushkina et al. 2014; Bouckaert et al. 2019; Zhang and Drummond 2020; Douglas et al. 2021), and conditioning on the root and rho parameters. Specifically, we filtered the autosomal UCE dataset for the most clock-like loci that also had low RF distances between gene and species trees using modified R scripts from a prior study (Musher et al. 2019). The details of these analyses, along with our justification of priors, node constraints, and fossil calibrations can be found in the Supplementary Materials (Alvarenga 1983; Bertelli and Chiappe 2005; Almeida et al. 2022). We also modeled speciation and extinction rates using the R-package ‘TESS’ (Höhna et al. 2016) (see Supplementary Materials).

### Evaluation of gene tree discordance

Gene tree discordance arises from multiple sources, including biological processes like ILS and introgression and gene tree estimation error. To examine the drivers of gene tree discordance and thereby better interpret phylogenetic results, we first used R v4.3.1 (R Core Team 2023) to assess the relationship between the number of parsimony informative sites (PIS) at each locus and and gene tree discordance. PIS were defined as variable sites in the alignment wherein each variant nucleotide is represented by at least two samples. We then measured the RF distances (Robinson and Foulds 1981) between each collapsed gene tree and the inferred species tree for the 100% complete UCE dataset only (an examination of the remaining datasets can be found in the Supplemental Materials), and used RF as a measure of gene tree discordance. RF distances were quantified using the ‘RF.dist’ function in phangorn v2.11.1 (Schliep 2011). To estimate RF distances, we compared the gene trees from each dataset to their corresponding MSC tree. To test for differences in gene tree discordance, as measured by RF distance, between autosomal and Z-linked UCE’s, we used Kruskal-Wallis tests in base R (R Core Team 2023). We then ran generalized linear models using PIS, a metric of information content, as the independent variable and RF distance as the dependent variable for the 100% complete UCE dataset. To characterize the relationship between information content and gene tree topology, we compared AIC values for linear and logarithmic fits for each regression and chose the model with the lowest AIC for each dataset. This enabled us to examine the potential impact of stochastic error (error introduced by insufficient data) on gene tree discordance, as the R-squared from this analysis measures the proportion of variation in gene tree topology (RF distance) that is explained by alignment information content (PIS). If stochastic error is absent, the slope should not significantly differ from zero and/or R^2^ should be low. From this, we could also see if Z-linked markers behaved differently from autosomal markers, with the expectation that Z-linked UCE’s should have reduced levels of ILS and introgression relative to autosomal markers, and thus reduced gene-tree-to-species-tree discordance as measured using RF distance. Specifically, we expect reduced RF distances for Z-linked gene trees relative to autosomal gene trees given comparable levels of phylogenetic information content (PIS). In other words, we expect that, while accounting for potential impacts of stochastic error driven by insufficient data in the alignments, RF distances will be reduced for Z-linked trees.

Because we found conflicting phylogenetic relationships between datasets for a sub-clade in the genus *Crypturellus* (Clade A, see Results), we assessed the extent to which this gene tree discordance could be explained by ILS due to rapid divergences or introgression between non-sister taxa. To do so, we first filtered all UCE alignments to only include one member of each species within Clade A (10 total plus one outgroup, as above for sliding window datasets), and then re-filtered for 100% completeness (including fewer samples resulted in more alignment with complete representation; complete sampling is a prerequisite for the quartet-based ILS analysis outlined below). We then reran IQ-TREE, using the same model selection, bootstrapping, and node collapsing as outlined above, to produce gene trees for the Clade A datasets. We then evaluated the impacts of ILS and introgression on the relationships in Clade A in multiple ways. Analyses found in the Supplemental Materials document, consistent with examples in the literature (Reddy et al. 2017; Stiller et al. 2024), suggested that BUSCO’s might be enriched for systematic error, so we therefore only applied these analyses to the UCE dataset.

First, we measured the relative quartet frequencies for two internal branches in Clade A on both Z-linked and Autosomal UCE’s. Quartets are unrooted four-taxon statements for which only three alternative topologies exist. The MSC model assumes that all gene tree discordance arises via ILS (i.e., no introgression, no recombination, etc.), and it makes predictions about the relative frequencies of alternative quartet topologies for all internal branches in a phylogeny. Specifically, the MSC predicts one majority topology consistent with the species tree at a relative frequency of >⅓ and two minority topologies of equal frequency at <⅓ (Pamilo and Nei 1988). As ILS increases, the frequencies of all three alternative quartets approach the ⅓ threshold. Deviations from these expectations (i.e., minority topologies are not equivalent in frequency) suggest that additional processes may be influencing gene tree discordance. For this method, we utilized the R package ‘MSCquartets’ (Rhodes et al. 2021; Allman et al. 2023). Because ILS is expected at relatively short internal branches, we evaluated the relative frequencies of alternative quartet topologies at two internal branches of the difficult-to-resolve clade (Clade A). To test for deviations from MSC predictions, we used a multinomial test in the R-package EMT (Menzel 2024) using the Monte Carlo approach to perform 10,000,000 simulated withdrawals based on expected quartet frequencies. We calculated the expected quartet frequencies for these branches using the ‘expectedCFs’ function in MSCQuartets on the UCE MSC species tree, which was congruent with the most common Clade A topology among datasets (see Results and Supplementary Materials).

To test for putative introgression among taxa in Clade A, we first estimated f_DM_ for 100 kb windows across the whole genome. F_DM_ is an f-branch statistic derived from the ABBA-BABA test (D-statistic) that can quantify the magnitude of introgression assuming a species tree topology. To do so, we first obtained genome-wide single nucleotide polymorphism (SNP) data for one member of each species within Clade A (*C. c. goldmani*, *C. erythropus, C. atrocapillus, C. duidae, C. kerriae, C. noctivagus, C. boucardi, C. strigulosus, C. transfasciatus, and C. u. undulatus*) plus one outgroup, *C. variegatus*. We again mapped cleaned reads for these samples to a reference genome, following the same protocol as outlined above, but this time using the pseudo-chromosome assembly from the Clade A ingroup with a relatively complete genome assembly, *C. u. undulatus* (LSUMNS 34614), as the same reference genome for all samples. We then used bcftools to call SNPs for all variants across the genome of each species. Finally, we filtered this merged vcf file using vcftools v1.16 (Danecek et al. 2011) to obtain SNPs with a minimum allele frequency = 0.05, maximum missing data = 0.5, minimum quality = 30, minimum depth = 10, and maximum depth = 100. Using these SNP data, we then estimated F_DM_ for 100 kb non-overlapping windows across the genome using the ‘ABBABABAwindows.py’ script located at https://github.com/simonhmartin/genomics_general#trees-for-sliding-windows (Martin et al. 2014). We specifically tested for introgression at branches whose phylogenetic positions differed among analyses and datasets. Because our datasets and phylogenetic analyses did not agree on a single most probable topology for Clade A, we ran this script four times, each time assuming one of the three hypothesized topologies (T1, T2, or T3; T4 had low support and was considered unresolved; see Results, Figure 2) as our guide tree.

Finally, we estimated the species tree topology while accounting for gene flow using a Bayesian approach implemented in PhyloNet v3.8.0 (the ‘MCMC_GT’ algorithm) (Wen et al. 2016). This method employs a reverse-jump Markov Chain Monte Carlo (rjMCMC) to sample the posterior distribution of phylogenetic networks under a multi-species network coalescent (MSNC) model using rooted gene trees as input. The MSNC is similar to the MSC, but relaxes the assumption of no introgression by modeling genome evolution as a network rather than bifurcating phylogeny. We ran the analysis twice, again using the gene trees with collapsed low-support nodes from the Z-linked and Autosomal Clade A UCE’s varying the maximum number of reticulate nodes in each run to assess variability in the results. We ran PhyloNet 11 times for both the autosomal and Z-linked datasets restricted to Clade A, once each for m-values between zero and ten (m defines the maximum number of reticulate nodes allowed in the MSNC model). We ran the rjMCMC for 5 x 10^7^ generations with a burn-in of 10%, using the pseudo-likelihood calculation. Because likelihood scores tended to increase with each successive increase in m-value, we chose the optimal m-value using a breakpoint analysis in the R package ‘segmented’ (Muggeo et al. 2014). The optimal m-value was chosen by fitting a segmented linear model to our likelihoods for each m-value. This allowed us to identify breakpoints in the slope of the regression, where increases in m-value resulted in diminishing gains in likelihood.

## Results

### Phylogenomics of Tinamous

The concatenated and MSC phylogenies for tinamous were well-supported and broadly congruent across datasets (Figures 1,2, S1–S21). The 75% and 100% complete datasets were congruent except for in the concatenated UCE results, implying that missing data had limited impact on phylogenetic reconstruction. However, there was slight variation in bootstrap and gCF’s support values, with 75% complete datasets tending to have higher support (Figure 2). The phylogenies from all datasets were congruent except for relationships within a single clade containing ten species of *Crypturellus,* we refer to as Clade A (Figure 2). The placement of *C. atrocapillus* was most unstable but was often recovered as sister to a clade containing *C. boucardi, C. kerriae, C. erythropus, C. duidae,* and *C. strigulosus* (Figure 2 “T1”). Overall, we recovered four different topologies for this clade based on the primary datasets (Figure 2: T1– T4). However, the Z-linked UCE dataset had low support for the placement of *C. atrocapillus* in the ASTRAL-III analysis (posterior probability = 0.46), and therefore failed to resolve the relationships of Clade A.

**Figure 1:**
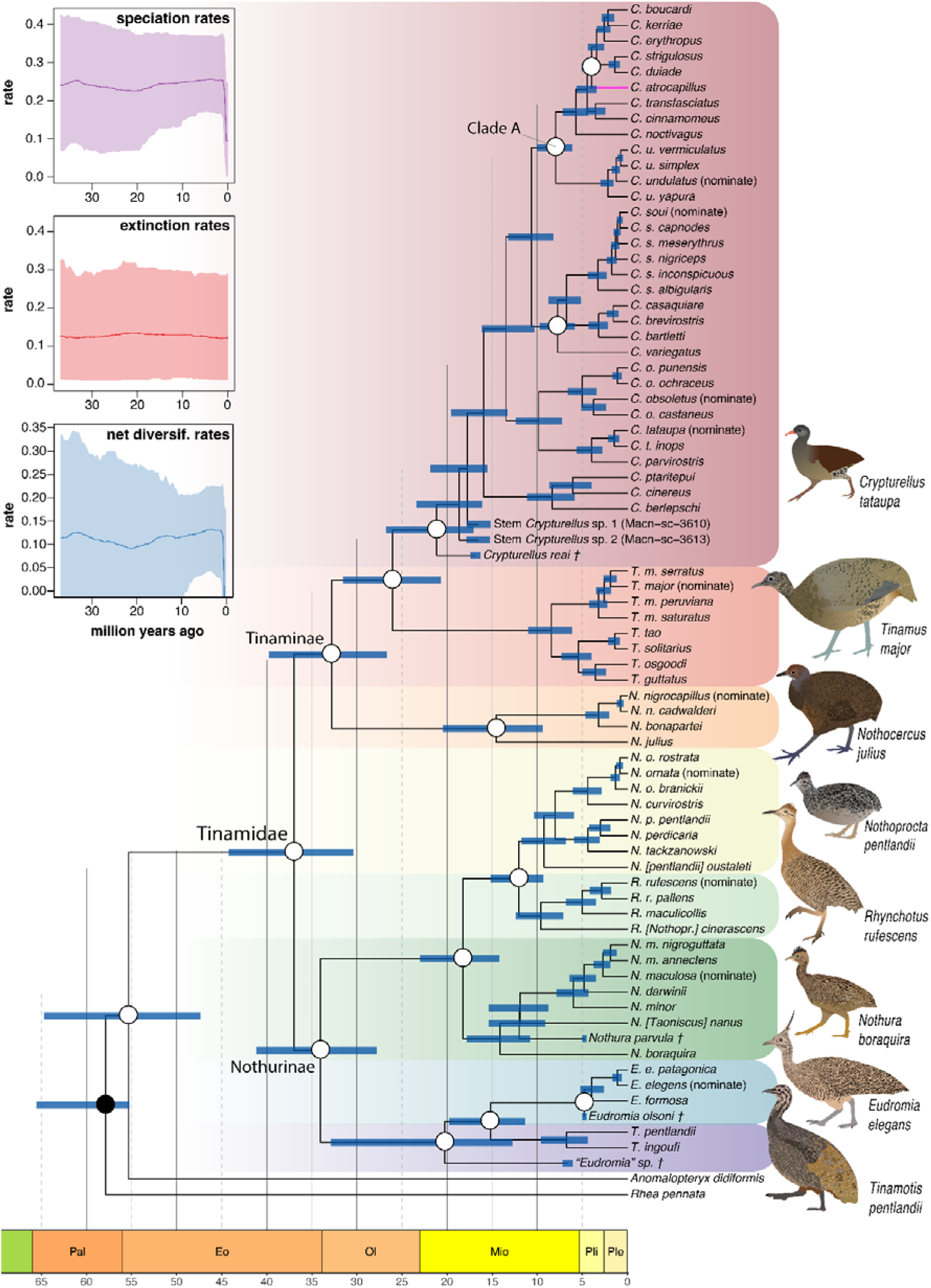
Time-calibrated Phylogeny of tinamous based on Autosomal UCEs with 1,000 bp of flanking sequence. Blue node bars represent 95% Highest Posterior Densities for node ages. White circles denote nodes constrained for monophyly based on maximum likelihood and MSC results. Tinamou illustrations by TAC.

**Figure 2:**
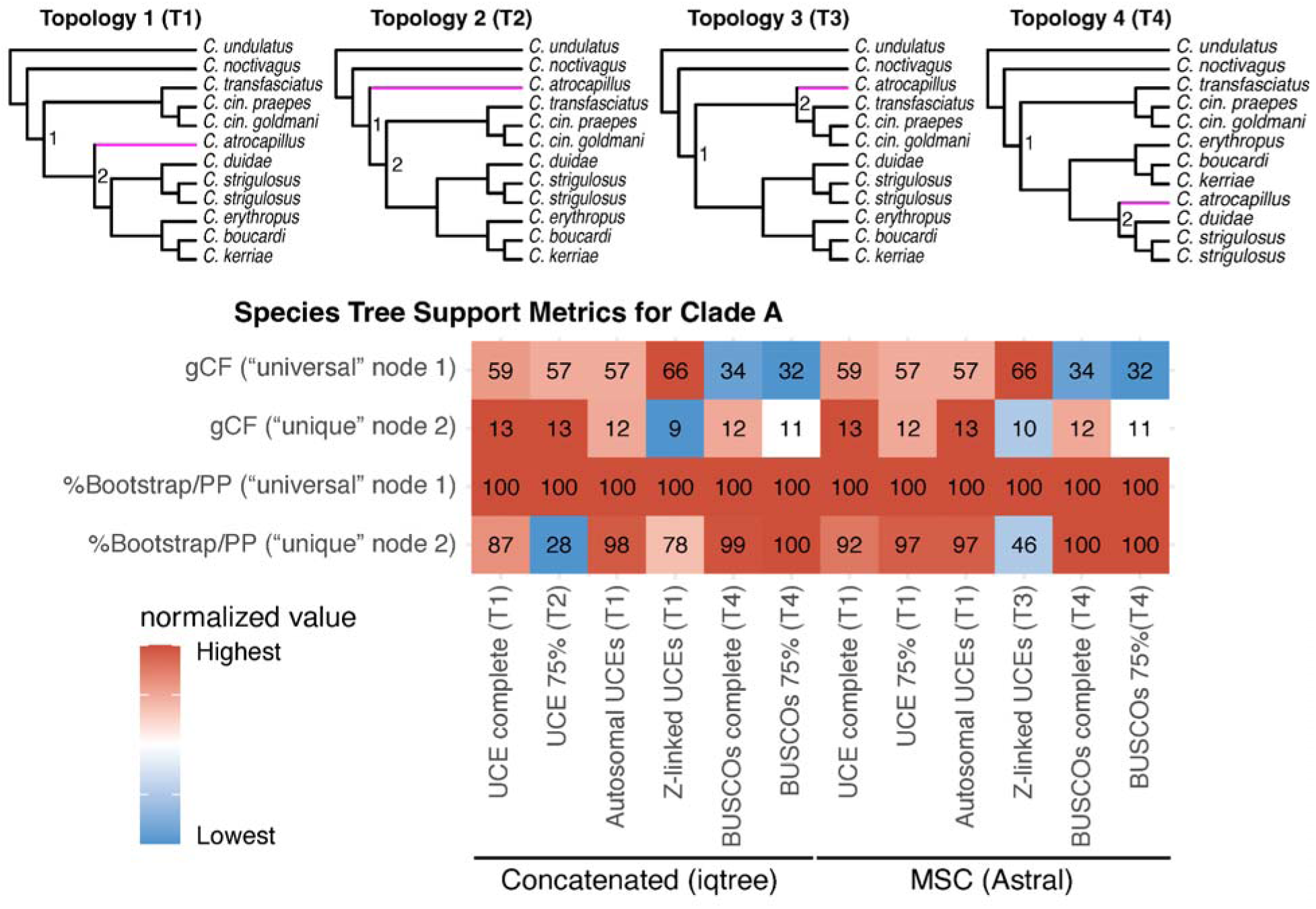
Phylogenetic topologies for a difficult-to-resolve clade (Clade A) of *Crypturellus* tinamous. The phylogenetic placement of *C. atrocapillus* (pink branch) was discordant among datasets and species tree approaches. The top four topologies are shown at the top (T1–T4). The heatmap at the bottom shows node support and/or concordance for each dataset and phylogenetic approach (concatenation and MSC). From top to bottom, these are (1) gene concordance factors (gCF) for a “universal” node shared among all three topologies, (2) gCF for a “unique” node in each topology associated with the position of *C. atrocapillus*, (3) bootstrap (IQ-TREE) or posterior probability (ASTRAL-III) for the “universal” node, and (4) bootstrap or PP for the “unique” node. The “universal” (node 1) and “unique” (node 2) nodes on each of the four topologies are marked on trees at the top of the figure. The numbers on the heatmap denote the rounded values, and the colors correspond to normalized values, such that the darkest red in each row is the highest value and the darkest blue is the lowest for that row. PP and bootstrap values are both shown as percentages for easy comparison. Results for additional datasets can be found in the Supplemental Materials.

Our results were broadly consistent with recognized taxonomic classifications. We recovered monophyletic subfamilies Nothurinae and Tinaminae and most genera and species were also monophyletic. However, two genera were non-monophyletic: the monotypic genus *Taoniscus* was embedded within *Nothura*, and *Rhynchotus* was embedded within *Nothoprocta*, rendering *Nothura* and *Nothoprocta* paraphyletic, respectively. We also recovered *Nothoprocta pentlandii* as polyphyletic; one specimen from Bolivia (voucher LSUMZ 123403) was sister to *N. perdicaria*, whereas the remaining samples from Peru formed a clade that was sister to all remaining *Nothoprocta*. All other species that were represented by multiple samples were recovered as monophyletic.

### The tempo of tinamou diversification

Our tip-dated phylogeny revealed relatively constant diversification of tinamous, which diverged from Moas roughly 55mya (Figure 1). The crown divergence of tinamous, separating forest (Tinaminae) from non-forest (Nothurinae) lineages, occurred roughly 37mya. A list of divergence times for genera and major clades can be found in Table S2. Speciation and extinction rates were stable through time, without any significant rate changes (Figure S25). We recovered a high rate of molecular evolution for the Tinamidae stem lineage (0.0034 substitutions/million years) compared to the mean for all lineages (ORCucldMean = 0.0019; 95% HPD = 0.0016–0.0023).

### Gene tree discordance arises from stochastic error, ILS, and introgression

We examined gene tree discordance for each dataset using RF distances between gene and species trees. To quantify the relationship between alignment information content, as measured using PIS, and gene tree discordance, we modeled RF distances as a function of the number of PIS per locus using a generalized linear model (Figure 3A & S23). This model was overall consistent with an expected negative association between PIS and RF distance, with lower RF distances in gene trees built from more informative alignments. This negative association (R^2^=0.29, *P*<0.0001) indicated that nearly one third of the variation in RF distance in UCE gene trees was explained by the number of PIS. This relationship was also logarithmic, suggesting that the effect of PIS on gene tree discordance was greatest for the least informative alignments but this effect quickly declined. When comparing autosomal and Z-linked UCE’s in this context, Z-linked UCE’s primarily fell below the model trendline, suggesting that for the same values of PIS, Z-linked UCE’s more closely resembled the species tree. Indeed, overall, Z-linked UCE gene trees averaged fewer differences from the species tree (mean RF distance between gene and species tree = 32.23.68 ± 9.67) than autosomal gene trees (mean=38.58 ± 9.67), and this difference was significant (*X*^2^=26.65, df=1, P<0.0001). Taken together, these findings suggest lower rates of ILS and/or introgression on the Z-chromosome, even when accounting for potential stochastic error.

**Figure 3:**
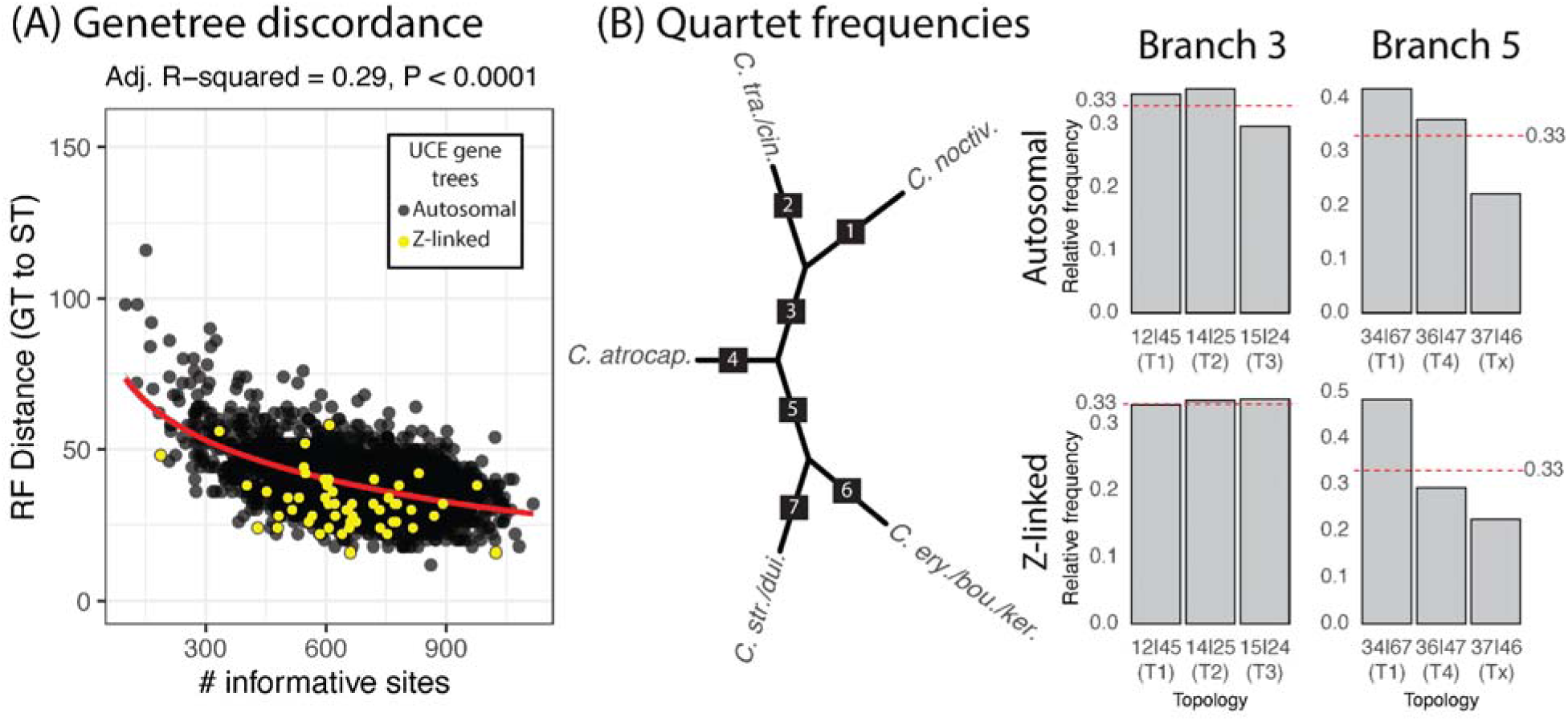
Gene tree discordance for autosomal and sex-linked UCEs. (A) Linear regression model exploring the relationship between information content (# parsimony informative sites) and gene tree discordance (RF distance between gene and species tree). The negative correlation suggests that alignments with less information content result in larger RF distances, but that Z-linked UCE’s (yellow points), fall below the trendline and thus have reduced RF distances given the same *x-*axis values. (B) Relative quartet frequencies for alternative unrooted topologies in Clade A. T1–T4 correspond to quartets consistent with topologies shown in Figure 2. Numbers on the branches of the unrooted network at the left coincide with numbers at the x-axis of each bar plot. For example, 12|45 denotes an unrooted quartet consistent with Topology 1 (T1; Figure 2) wherein *C. noctivagus* (branch 1) is sister to *C. transfasciatus* + *C. cinnamomeus* (branch 2) and *C. atrocapillus* (branch 4) is sister to branch 5. “Tx” refers to a quartet that does not match one of the four primary topologies in Figure 2.

We found evidence of ILS and deviations from the MSC model using the quartet approach (Figure 3B). For one branch (branch 3), we recovered frequencies for three alternative quartets that approached the ⅓ threshold when using autosomal UCE’s, implying ILS has promoted phylogenetic discordance at these branches. However, two of the alternative quartets exceeded the ⅓ threshold and were roughly equal in frequency, indicating a deviation from the MSC. When assessing quartet frequencies for Z-linked gene trees at branch 3, the results closely matched expectations of ILS, wherein the relative frequencies of the three alternative quartets were approximately equal to the ⅓ threshold, and we did not reject a null model where Z-linked quartets had equivalent frequencies based on the multinomial test (P=0.88). For another branch (branch 5), we found weaker evidence for ILS, with a single majority topology (34|67) and two minority topologies. For both autosomal and Z-linked gene trees, we also found deviations from pure ILS, as the two minority topologies were not equal in frequency. However, we again found that Z-linked gene trees matched expectations of reduced ILS on the sex chromosomes, with a majority topology becoming more frequent in the Z-linked quartet analysis (i.e., less gene tree discordance). In all cases, quartet frequencies differed from MSC expectations given the UCE ASTRAL-III species tree (P<0.0001).

Our genome-wide analysis of f_DM_ for 100 kb sliding windows provided contrasting signals for introgression across the genome, but identified non-zero signals of introgression across most windows of the genome (Figure 4A). When assuming T1 as the phylogenetic background in the model, we found slight negative values of f_dM_ across autosomes indicating introgression between the *C. cinnamomeus/transfasciatus* branch and the *C. strigulosus/duidae/erythropus/kerriae/boucardi* branch (P1 and P3 in this model). These values of f_dM_ only slightly deviated from zero (mean autosomal f_dM_ = −0.021 ± 0.00022), whereas the analysis recovered very strong signals of introgression for many windows along much of the Z-chromosome (mean Z-linked f_dM_ = −0.117 ± 0.0052), and this difference was significant (*X^2^* = 311.21, *df* = 1, *P* < 0.0001). When assuming T2 as the phylogenetic background, we saw the opposite pattern; autosomes again showed consistent negative values of f_dM_ indicating introgression between *C. atrocapillus* and the *C. strigulosus/duidae/erythropus/kerriae/boucardi* branch (mean autosomal f_dM_ = −0.027 ± 0.00029), but the windows on the Z-chromosome averaged much closer to zero (mean Z-linked f_dM_ = −0.010 ± 0.00098). The difference between mean autosomal and Z-linked introgression was significant (*X^2^*= 293.49, *df* = 1, *P* < 0.0001). When assuming T3 as the phylogenetic background, we again found evidence of genome-wide introgression, this time mostly positive values indicating introgression between *C. atrocapillus* and the *C. strigulosus/duidae* branch. There were several obvious negative peaks across the genome in this scenario, including much of the Z-chromosome (mean autosomal *f_dM_* = 0.027 ± 0.00037; mean Z-linked *f_dM_* = −0.022 ± 0.0019), indicating introgression between *C. atrocapillus* and the *C. erythropus/kerriae/boucardi* branch.

**Figure 4:**
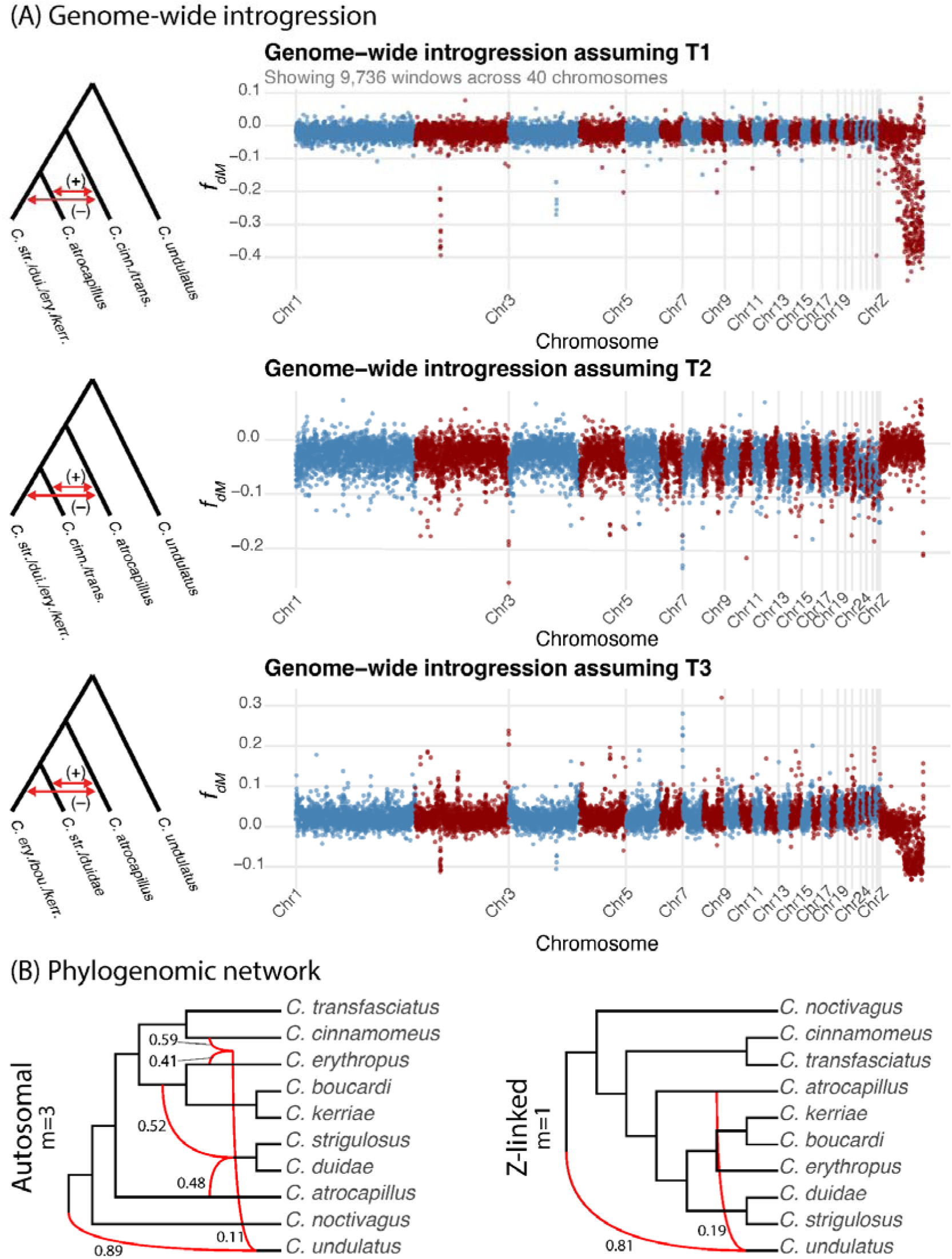
Analysis of genome-wide introgression. (A) Introgression (f_dM_) mapping for 100 kb non-overlapping windows across the genome assuming each of the three well supported topologies (Figure 2, T1–T3). The trees at the left of each plot show the assumed background topology and red arrows indicate positive or negative values of f_dM_. Tips on the trees show taxa assigned to each population in the introgression model (from left to right: P1, P2, P3, and P4 assigned populations in the model). (B) The results from the multi-species network coalescent model in PhyloNet. The Z-linked tree was weakly supported.

As a final test of the Clade A topology and impacts of gene flow, our phylogenomic network analysis revealed multiple nodes involved in reticulation that may explain gene tree discordance in Clade A (Figure 4B). A breakpoint analysis indicated that the optimal m-value (number of reticulate nodes) was m=3 for autosomal UCE’s and was m=1 for Z-linked UCE’s (Figure S26); thus, we recovered fewer introgression events for the Z-linked than autosomal UCE’s. For autosomal UCE’s, we found a network with a reticulate node involving *C. atrocapillus* (posterior probability = 1.0), suggesting that uncertainty around the placement of this species could conceivably be driven by historical introgression. For the Z-linked loci, we recovered a network most closely resembling T1 with a signal of introgression between *C. undulatus* and *C. atrocapillus,* but this specific network topology was not supported (posterior probability = 0.46).

## Discussion

Here we used whole-genome sequencing to explore the sources of phylogenomic conflict and reconstruct the species-level phylogenetic history of tinamous, a relatively old Neotropical avian group of broad interest in evolutionary biology (Sackton et al. 2019; Li et al. 2023). Our results based on different analytical approaches (concatenated and MSC methods) and datasets (UCE’s and BUSCO’s) were largely congruent. The topology of only a single clade in the genus *Crypturellus* (Clade A), which contained multiple successive short internodal branches, varied among approaches. These results indicate that species tree estimation in this group was, in general, robust, but that gene tree estimation error, ILS, and introgression may have been key drivers of gene tree discordance that hampered phylogenetic inference for one recalcitrant node in the genus *Crypturellus* (in Clade A).

Indeed, both ILS and introgression likely drove uncertainty in the position of *C. atrocapillus*, the most unstable node in the phylogeny, which had relatively low concordance factors and reduced node support in many datasets (Figure 2). The relative quartet frequencies for this node closely approached the ⅓ threshold predicted by high ILS, especially from Z-chromosome alignments (Figure 3B). However, given that we could not eliminate the impacts of stochastic error, it remains uncertain how much of the gene tree discordance described in the quartet analysis was associated with ILS versus gene tree estimation error. In addition to ILS and introgression, rapid divergence events can be difficult to reconstruct because the short timeframe also means there is limited phylogenetic signal for those divergence events within individual gene trees (Leaché et al. 2015; Mclean et al. 2019). Still, multiple methods pointed towards genome-wide introgression in this clade, potentially depleting signal for the species tree across much of the genome and further complicating phylogenetic reconstruction in Clade A. Additional analysis and intraspecific sampling may be necessary to determine the precise phylogenetic relationships of taxa in Clade A (Arcila et al. 2017; Shen et al. 2017; Remsen et al. 2024).

### Phylogeny and taxonomy of the Tinamous

Our phylogenies based on multiple datasets all resulted in highly similar and well-supported relationships among tinamou species. For example, past studies, based primarily on morphological data or few molecular loci, recovered high uncertainty in the interrelationships of *Crypturellus* species (Bertelli et al. 2014; Almeida et al. 2022), but our reconstructions for this genus were mostly robust. Moreover, concatenation and MSC approaches largely agreed on the overarching relationships at both shallow and deeper timescales. Importantly, our phylogenetic results imply that multiple taxonomic changes are necessary to appropriately classify taxa within Tinamidae. First, *Nothoprocta cinerascens* was recovered as the sister to the genus *Rhychotus* (Spix 1825). *Rhynchotus* (Spix 1825) has nomenclatural priority over *Nothoprocta* (Sclater and Salvin 1873), and we therefore suggest moving *N. cinerascens* to the genus *Rhynchotus* (Bertelli and Porzecanski 2004; Valqui 2009). Second, *Taoniscus nanus* was nested within the genus *Nothura*, and thus, *Taoniscus* (Gloger 1842) is a junior synonym of *Nothura* (Wagler 1827). We therefore suggest transferring *T. nanus* to *Nothura.* Third, *C. duidae*, often considered a close relative of *C. noctivagus* or *C. erythropus* (Remsen et al. 2024), was found not to have close affinities to either of these; instead, it was consistently placed as sister to *C. strigulosus.* Finally, we found evidence for what is likely an unrecognized species of *Nothoprocta* (Valqui 2009); specimens ascribed to *N. pentlandii* were polyphyletic, with a specimen of the nominate subspecies from Bolivia recovered as sister to *N. perdicaria*, and multiple specimens from Peru recovered as a clade sister to all other *Nothoprocta.* Thus, there appear to be at least two species-level taxa within *N. pentlandii*, and more work is needed to properly circumscribe the taxa in this species complex.

### Tip-dating enables approximation of the relationships of fossils even when phylogenetic data are equivocal

Our estimates of divergence dates for tinamous were broadly consistent with a recent study (Almeida et al. 2022). Although we used many of the same fossils for calibration, Almeida et al. (2022) used node calibrations. Here we used tip-dating, which explicitly accounts for uncertainty in fossil placement, enabling us to include additional crown fossils never before included in a dating analysis. Although the placement of some fossils in our dataset have been recovered with strong support in previous studies, others have not (Bertelli et al. 2014; Almeida et al. 2022). For example, prior work has found that an extinct *Eudromia-*like species is sister to the clade containing extant *Eudromia* and *Tinamotis,* and two undescribed fossil fragments (MACN-SC-3610 and MACN-SC-3613) belong to stem *Crypturellus*. On the other hand, the placement of the extinct *C. reai*, was equivocal in morphological phylogenies despite sharing clear synapomorphies with extant *Crypturellus*. Our analysis allowed the placement of *C. reai* to vary between crown and stem *Crypturellus*, and showed that this species either belongs to an early crown lineage or late stem lineage, but is not likely closely related to any extant species (Figure 1). For example, the posterior probability of crown *Crypturellus* in our tip-dated tree was 0.4, indicating a possibility that *C. reai* could be nested within the crown. However, all other nodes in this clade were strongly supported as monophyletic (PP≥0.98; supplemental tree files).

In line with other macroevolutionary studies for Neotropical birds, we found relatively constant rates of diversification for tinamous (Patel et al. 2011; d’Horta et al. 2013; Harvey et al. 2020). Our results agree with those of Almeida et al. (2022) that tinamous diverged from their sister group, the extinct moas, during the Paleocene or early Eocene with crown tinamous diverging into forest (Tinaminae) and open-country (Nothurinae) clades during late Eocene or early Oligocene (Figure 1; Table S6). All extant genera of tinamous arose in the mid-to-late Miocene, except *Eudromia* (Late Miocene to Pliocene) and *Crypturellus* (early-to-mid Miocene). Using our unique approach, we also recapitulated the high substitution rate of the stem tinamid lineage, one of the highest in modern birds (Berv and Field 2018), a finding that remains unexplained.

### The drivers of gene tree discordance in Clade A include pervasive signals of introgression

We examined orthologous data harvested from whole genome assemblies for the potential impacts of stochastic error to better inform our tests of ILS and introgression. We found that, as expected, as parsimony informative sites (PIS) are added to alignments, gene-tree-to-species-tree discordance is reduced. This relationship is well known (Meiklejohn et al. 2016; Burbrink et al. 2020), but confirming it for our dataset allowed us to understand baseline levels of potential stochastic error in the data (i.e., 29% of the variance in RF distances is explained by alignment information content), and thus evaluate levels of ILS in light of this relationship. Z-linked gene trees fell below the trendline for the PIS x RF distance regression model, implying that for the same number of PIS as autosomal alignments, Z-linked alignments had lower RF distances. We argue that this is likely because Z-linked UCE’s should have reduced rates of ILS, though disentangling stochastic and other forms of error from legitimate biological signals of ILS in these gene tree summarization methods is admittedly difficult.

In Clade A, the placement of *C. atrocapillus* varied among datasets and analytical approaches, but one topology was most common (T1). This topology was recovered for most UCE trees as well as the concatenated Z-linked UCE tree (Figure 2). Sex-linked markers are expected to experience reduced rates of ILS and faster coalescent times than autosomes due to smaller effective population size, as well as reduced rates of recombination and introgression (Charlesworth et al. 1987; Meisel and Connallon 2013; Li et al. 2019; Rivas-González et al. 2023; Musher et al. 2024; Burbrink et al. 2025). These expectations are consistent with our findings that (1) as laid out above, RF distances between gene and species trees were lower for Z-linked UCE’s than for autosomal UCE’s given comparable values of PIS, suggesting reduced ILS on the Z-chromosome, (2) the quartet frequencies of alternative topologies for relatively short internal branch lengths more closely matched MSC model expectations for Z-linked than autosomal UCE’s, suggesting reduced introgression on the Z-chromosome, and (3) the optimal number of reticulation events was lower for Z-linked phylogenetic network than for the autosomal network, though this last observation could be driven, in part, by the large difference in the number of alignments obtained for each dataset (3,942 autosomal UCE’s vs. 200 Z-linked UCE’s). Nevertheless, the gene trees from the Z-linked dataset failed to resolve the species tree and network for Clade A on their own. It thus seems likely that given the small number of Z-linked genes analyzed, and rapid divergence at the *C. atrocapillus* node, a larger Z-linked dataset may be needed to resolve the relationships in Clade A. Future work might generate Z-chromosome alignments to obtain more data that could help resolve Clade A.

Interestingly, our analysis of introgression using f_dM_, which does not rely on gene tree estimation, may provide some insight into the species tree of Clade A. When assuming T1, the most commonly recovered topology in our study, as the species tree, we found low levels of introgression for most sliding windows across the autosomes but significant peaks of introgression (negative f_DM_ values) along those on the Z-chromosome, a finding that is opposed to theoretical expectations that the Z-chromosome should have reduced introgression (Burbrink et al. 2025). This result is striking and could be interpreted to mean that T1 is the wrong topology despite being the obvious candidate for the species tree. Contrastingly, when assuming T2 as the species tree, we recovered levels of introgression (negative f_DM_ values) that were consistent with such expectations: pervasive signals of introgression across all the autosomes, with values closer to zero on the Z-chromosome. These results raise the possibility that T2, despite only being recovered in one analysis, might be the correct species tree, but the signal for T2 has been depleted in the genome by the overwriting impacts of pervasive autosomal introgression (Fontaine et al. 2015). T2 was also the most frequent autosomal quartet for branch 3 (Figure 3B). Pervasive introgression between *C. atrocapillus* and the ancestor of *C. erythropus, C. boucardi*, and *C. kerriae* could explain both the difficulty in determining the phylogenetic placement of *C. atrocapillus*, while also explaining why the T1 was the most commonly recovered topology (i.e., it may be a signal of introgression). Indeed, our phylogenomic network from autosomes, which estimated the phylogenetic history while accounting for introgression, recovered a network involving reticulation between *C. atrocapillus*, the *C. strigulosus* + *duidae* clade, and the *C. erythropus + boucardi + kerriae* clade. However, we would argue that more work is needed to corroborate this hypothesis. Instead, we conservatively suggest that the phylogeny of Clade A remains uncertain, with evidence that rapid divergence and introgression have promoted this phylogenetic uncertainty.

## Acknowledgements

We thank Alvaro Hernandez and Chris Wright at the University of Illinois Roy J. Carver Biotechnology Center for assistance with Illumina sequencing. We thank Kimberly K. O. Walden for assistance in depositing Illumina reads to NCBI SRA. We also thank Jorge Doña for providing R code to assist with breakpoint analysis. Dan Lane, Steve Cardiff, and Christopher Witt helped confirm voucher identities for multiple samples. We thank B. T. Smith, J. Cracraft, P. Sweet, T. Trombone, S. Katanova (AMNH), J. Bates, S. Hackett, and B. Marks (FMNH), F. Sheldon and D. Dittman (LSUMNS), H.F. James and B. Schmidt (USNM), and N. Rice (ANSP) for loaning material that was crucial for this work. Finally, we thank the members of ANSP Ornithology Department (A. Del Grosso, E. Griffith, K. Kuabara, and J. Merwin) for comments on early versions of the manuscript. Comments from S. Ruane, Isaac Overcast, and five anonymous referees greatly improved this work. This study was funded by NSF grant DEB-1855812 to JDW and KPJ. TAC was partially supported by NSF grant DEB-2203228.

## Data Availability

All raw reads are available on the NCBI Sequence Read Archive (project and sample accession numbers can be found in Table S1). All the original data and scripts necessary to reproduce the analyses reported in this study can be accessed through LJM’s github page: https://github.com/lukemusher/Tinamou-phylogenomics.git.

