## Supplemental Materials for "Whole-genomes illuminate the drivers of gene tree discordance and the tempo of tinamou diversification (Aves: Tinamidae)"

**Overview**

Herein, we present supplementary Methods, Results, Discussion, Figures, and Tables. We expand on and discuss analyses that scrutinize the covariates of gene tree discordance, with the objective of examining levels of stochastic error in our data.

*Background on gene tree estimation error*

High throughput datasets can contain biased gene alignments that artificially increase gene tree discordance, mimicking the effects of ILS or introgression, and thus biasing phylogenetic estimation under standard approaches (Doyle et al. 2015; Smith et al. 2018; Zhao et al. 2023). For instance, multi-species coalescent (MSC) methods that utilize gene trees as input may fail if gene tree discordance is driven by estimation error rather than coalescent variation (Xi et al. 2015; Meiklejohn et al. 2016; Springer and Gatesy 2016; Simmons and Gatesy 2021). Shorter alignments with relatively few parsimony informative sites (PIS) may result in poorly resolved gene trees due to a lack of phylogenetic signal in the data, a problem referred to as stochastic error (Xi et al. 2015; Meiklejohn et al. 2016; Kapli et al. 2021). Conversely, utilizing alignments of long loci that contain many PIS is expected to reduce stochastic error. Some studies have shown a negative correlation between the number of PIS and discordance between gene and species trees (Burbrink et al. 2020), implying that as PIS are added to alignments, stochastic error is reduced. On the other hand, increasing alignment length also increases the probability of sampling multiple independently sorting coalescent genes (*c*-genes; genomic segments with distinct branching histories), which may range in size from tens of base pairs in length for deep divergences to a few thousand base pairs for recent divergences, and this is known to affect phylogenetic inference (Doyle 1995, 1997; Schierup and Hein 2000; Hobolth et al. 2007; Zhao et al. 2023). Another, more troublesome source of gene tree inaccuracy is systematic error, which is driven by data biases that cause model violations (e.g., GC content variation) (Rodríguez-Ezpeleta et al. 2007). Therefore, dissecting biological and artifactual sources of gene tree discordance to infer species trees more accurately is a major challenge in the age of phylogenomics, where more data does not automatically translate to improved inferences (Meiklejohn et al. 2016; Blom et al. 2017; Smith et al. 2023).

### Methods

*DNA extraction, library preparation, and sequencing*

For fresh tissues, we extracted genomic DNA using the MagAttract High Molecular Weight kit from Qiagen (Valencia, California). Toe pad extractions of historical study skins was carried out in a dedicated historical DNA extraction laboratory at the Academy of Natural Sciences of Drexel University to reduce contamination risk. Toe pad samples were first washed in a brief bath of EtOH to help remove superficial contaminants, and then soaked in $H_2O$ for three hours to hydrate the desiccated flesh for DNA lysis. Samples were then digested using 180 µL ATL and 30 µL proteinase K for each sample and incubated at 56º C overnight. DNA isolation was then performed using the QiaQuick spin columns and extraction kit from Qiagen (Valencia, CA).

Shotgun sequencing libraries were prepared for each extract using the Hyper library construction kit (Kapa Biosystems). These libraries were sequenced using 150 bp paired-end reads on an S4 lane of an Illumina NovaSeq 6000. These libraries were pooled and tagged with unique dual-end adaptors, and pooling consisted of between 13 and 18 samples per lane aimed to achieve at least 30X coverage of the nuclear genome. Adapters were trimmed and demultiplexed using bcl2fastq v.2.20. We deposited raw reads in the NCBI SRA (Table S1).

After mapping reads to reference genomes using BWA (see main text), we used '-doFasta 2' flag in ANGSD to ensure that the consensus nucleotide was written for each polymorphic site.

In an earlier draft of this manuscript we used "CryUnd" (NCBI GCA_013389825.1) as reference for several samples(Musher et al. 2024). During revisions we found that samples mapped to this genome were more likely to cluster with each other than with other more likely relatives in certain phylogenetic analyses. This led us to remap these samples to a different reference, *C. strigulosus* (LSUMNS B-9577), in the current version.

*Assessing genome completeness*

To assess genome quality, completeness, and redundancy for each assembly, we used Benchmarking Universal Single Copy Orthologs (BUSCO) version 5.3.0(Simão et al. 2015). BUSCO searches genome assemblies and identifies genes present in single copy using a database of known single-copy orthologs from a clade-specific database of genes. We used the 'aves_odb10' lineage dataset, which utilizes 8,338 genes in the chicken genome. We used the '-- augustus' flag to obtain nucleotide sequences for genes. This setting uses augustus version 3.2(Hoff et al. 2019) to annotate each assembly, and outputs full nucleotide gene sequences for all complete single-copy orthologs in fasta format. This step was necessary to obtain our coding gene dataset for phylogenomic analysis. We also used samtools to estimate mean and standard deviation of sequence coverage for each genome.

*Alignment trimming*

An earlier version of this study found that BUSCOs with more Parsimony Informative Sites did not significantly improve gene tree accuracy, implying a strong signal of systematic error in this dataset. We examined a handful of BUSCO alignments by eye and found long sequences unique to a small number of individuals in each alignment. We suspected that these were driven by misannotation in the BUSCO pipeline, and thus trimmed all loci using TrimAL to remove sites in each alignment with missing data for more than two samples. This seemed to alleviate the problem as we found a tighter correlation between the number of parsimony informative sites and RF distance between gene and species trees after trimming.

*Phyluce harvesting of UCEs from whole genomes and additional phylogenomic details*

We harvested UCEs three times to construct three UCE datasets that each varied in the length of the flanking region around the UCE core. These UCE datasets included 100 bp (henceforth UCE100 dataset), 300 bp (UCE300 dataset), and 1,000 bp (UCE1000 dataset; in the main text of this study, we discuss only this UCE set) of flanking region on both 5' and 3' ends of each conserved core. These three UCE sets are not subsets of one another, but independent extractions of the same UCE's with different flank sizes. To examine differences among sex-linked and autosomal markers, we also created two datasets of long UCEs (1,000 bp flanking regions), one of those that mapped to the Z-chromosome (an avian sex chromosome with idealized $N_e$ of 0.75 autosomal $N_e$) and one that mapped to autosomes (see Supplementary Materials).

To generate Z-linked and autosomal datasets, we first used the Phyluce scripts to harvest all UCEs from each pseudo-chromosome-level genome. We then made a list of UCE loci mapping to the Z-chromosome by using "grep" in UNIX to extract lines in the '.lastz' file containing UCEs on the Z-chromosome in each genome and "grep -v" to exclude lines on the Z-chromosome (autosomal dataset). This generated new '.lastz' files for each sample which were then used in the Phyluce pipeline.

For all datasets we used MAFFT v7.453 (Katoh and Standley 2013) to align orthologous UCE loci for downstream phylogenomic analysis and filtered these alignments into 75% complete (≥75% of samples represented in each gene alignment) and 100% complete (all samples represented in each gene alignment) alignments. However, for the autosomal and Z-chromosome UCE datasets, we only used 75% complete alignments to maximize alignment inclusion in these two datasets. This resulted in ten datasets (Table S4). All alignments were trimmed using TrimAL v.1.4 (Capella-Gutierrez et al. 2009) with a gap threshold of 97.5% to trim out potential BUSCO mis-annotations unique to a small number of samples (see Supplementary Materials). Each dataset was analyzed using the same phylogenetic methods as outlined in the main text, which resulted in species and gene trees for each of the ten datasets.

*Assessing ortholog quality and gene tree error*

To evaluate variation in phylogenetic information content and stochastic error among datasets, we estimated the number and proportion of parsimony informative sites (PIS) for each alignment within each dataset of 100% complete alignments (downstream analyses require full representation for all loci and gene trees). PIS were defined as variable sites in the alignment wherein each variant nucleotide is represented by at least two samples. To examine potential effects of systematic error, we also estimated GC content variation (variance in GC content among all sites in each alignment) for each alignment. To examine levels of gene tree discordance for each dataset, we then measured the Robinson-Foulds distances (RF distance) (Robinson and Foulds 1981) between each gene tree and the inferred species tree (using the inferred species tree for each dataset, rather than one species tree for all datasets). RF distances were quantified using the 'RF.dist' function in phangorn v2.11.1 (Schliep 2011). To test for differences in these measures among datasets, we used Kruskal-Wallis tests in base R (R Core Team 2023) followed by a post-hoc Dunn Test using the FSA package (Ogle et al. 2023). We expect that, if all gene tree discordance is associated with biological processes, the mean and variance in RF distances among datasets will be relatively constant. If the mean RF distance is higher for some datasets, it could mean that there is increased error in that dataset. Still, as alignment length increases, so does the number of PIS, but also the potential inclusion of multiple *c*-genes. So lower mean RF distances for one dataset could indicate convergence of gene tree topologies on the concatenated topology, rather than a reduction in error.

Gene tree discordance arises from multiple sources, including ILS, introgression, and potential stochastic or systematic error. One way to examine whether the measured discordance is biological or erroneous is to see how gene tree discordance covaries with alignment information content (an investigation of potential stochastic error) or GC content variation (an investigation of potential systematic error). Stochastic error arises because there is not enough information to reconstruct a gene history resulting in ambiguously or incorrectly resolved gene trees. If there is no impact of stochastic error, one would expect no statistical relationship between these PIS and gene tree discordance. Therefore, this analysis gives us a baseline understanding of rates of error and which datasets have reduced error. Moreover, it allows us to look for deviations from this relationship (e.g., between Z-linked and autosomal markers) to see if one of these datasets has lower rates of discordance (RF distance) given the same values of information content. Thus, to investigate covariation between gene tree discordance, PIS, and GC content variation, we then ran generalized linear models using information content (PIS) or GC content variation as the independent variables and RF distance. We compared AIC values for linear and logarithmic fits for each regression and chose the model with the lowest AIC for each dataset. These regression analyses were done once for each dataset, and then, to better investigate broader patterns of systematic error given both shorter and longer alignments, once with the three UCE datasets combined. To avoid pseudo-replication in the combined regression, we randomly removed replicated UCE's from the data-frame such that each UCE was only represented once (either by the UCE100, UCE300, or UCE1000 alignment).

To examine if differences between UCE and BUSCO regression results could be driven by systematic error, we explored these models again for BUSCO's after filtering the BUSCO dataset for GC content variation (by removing genes with GC content variation greater than two standard deviations from the mean) and for BUSCO's resembling UCE datasets (removing genes greater than two standard deviations above the mean proportion of PIS in the UCE1000 dataset and below the mean of the UCE100 dataset). The idea here is to attempt to replicate a dataset that looks like the combined UCEs in terms of PIS. Finally, we combined both previous filters to make a strictly filtered dataset.

*Estimating the tempo of tinamou diversification*

First, we filtered the autosomal dataset to include only a single member of each taxon (monotypic species or subspecies) and created a new 100% complete autosomal dataset. Next, we generated new gene trees for these alignments and adapted scripts from a prior study (Musher et al. 2019) to quantify the RF-distance between gene and species tree and a measure of clock-likeness for each alignment in the autosomal UCE dataset. Specifically, we estimated the likelihood ratio of clocklike to non-clocklike models for each gene; lower ratios indicate molecular evolution that is more clocklike. The protocol we followed was identical to that in Musher et al. (2019). After estimating likelihood ratios and RF-distances for all alignments and gene trees, we selected only those loci with RF distances below the mean distance, and with likelihood ratios that were two standard deviations below the mean ratio. This allowed us to include a relatively small number (22 in total) of genes with relatively low variance in molecular substitution rates among branches, but were relatively informative for phylogenetic reconstruction (Figure S27).

To estimate the timeframe of tinamou diversification, we used a fossilized birth-death model with six crown Tinamidae tip-fossil calibrations using an optimized relaxed clock and fossilized birth-death model in Beast v2.7.7 with the Sampled Ancestors and Optimized Relaxed Clock packages (Drummond et al. 2006; Gavryushkina et al. 2014; Bouckaert et al. 2019; Zhang and Drummond 2020; Douglas et al. 2021), and conditioning on the root and rho parameters. We then used the filtered autosomal UCE1000 alignments described above, generating lines of question marks in each alignment for six fossil taxa (see Table S2). We applied tip dates measured as years before the present using initial dates listed in Table S2, and uniform priors on tip ages using the upper and lower bounds of their estimated ages from the fossil record. We linked all clock and tree models but partitioned site models by locus (22 in all), with a GTR + G model for each partition. Default priors and settings were used except where specified in Table S3.

Because we used a relatively small number of DNA sequences, we constrained 14 nodes in the tree using MRCA monophyletic node constraints. First, we constrained all nodes with fossils of known position based on previous morphological work (Bertelli et al. 2014).

Second, after preliminary analysis without further node constraints, we constrained nodes that did not converge on the topologies known from the main phylogenetic analyses. The specific details of the monophyly constraints are as follows (1) crown Tinamidae, (2) crown Nothurinae, (3) crown Tinaminae, (4) "*Eudromia*" sp. + *Eudromia* + *Tinamotis,* (5) *Eudromia* + *Tinamotis*, (6) *Eudromia*, (7) *Nothoprocta* + *Rhynchotus* + *Nothura*, (8) *Nothoprocta* + *Rhynchotus*, (9) *Nothocercus*, (10) *Tinamus* + *Crypturellus*, (11) *Crypturellus* including fossils Macn-sc-3610 and Macn-sc-3613, (12) *Crypturellus soui, C. variegatus, C. bartletti, C. brevirostris,* + *C. casaquiare,* (13) *Crypturellus* Clade A, and (14) *C. boucardi, C. kerriae, C. erythropus, C. strigulosus,* + *C. duidae*. The final constraint was used to match topology T1, the most frequently recovered topology in our phylogenetic analyses of the most difficult node in the phylogeny. We ran the MCMC for 150,000,000 generations with sampling every 50,000 generations. The tree was summarized as a maximum clade credibility tree with median values for node heights, and a burn-in of 20% in TreeAnnotator v2.7.7(Bouckaert et al. 2019). Convergence was assessed using Tracer v1.7.1(Rambaut et al. 2018).

We also modeled speciation and extinction rates using the R-package 'TESS' (Höhna et al. 2016). TESS uses a rjMCMC algorithm to estimate the timing of speciation and extinction rate shifts under an episodic birth-death model. To do so, we pruned fossil tips and outgroups out of the beast tree, and ran the MCMC for $10^6$ generations, assuming 85% complete taxonomic sampling for the clade.

### Results

*Assembly metrics, completeness, and ortholog metrics*

We successfully assembled 68 of the 69 newly sequenced genomes and extracted BUSCO and UCE targets from these assemblies plus 10 publicly available tinamou and 2 publicly available outgroup whole genome assemblies (Table S1). Genome-wide sequence coverage varied within and among samples, but was generally high; the mean of average coverage across samples was 40.06x (standard deviation = 14.13x). Most genomes were also relatively complete, containing an average of 89.69% (standard deviation = 10.88%) of 8,338 complete single-copy BUSCO genes.

One sample (*C. soui poliocephalus* ANSP 187533) had relatively low coverage (mean coverage = 10.42x, standard deviation = 76.95x) and thus read-mapping resulted in a highly incomplete genome (Table S1). This sample was dropped from all subsequent analyses. Another genome, downloaded from NCBI (GCA 013389825) was labeled as *C. undulatus*, but did not cluster with other members of that species, instead clustering with *C. strigulosus* (most trees). We could not confirm the species identity of this sample because the voucher was unavailable or lost, but photographs of the headless voucher are consistent with *C. strigulosus*. Moreover, another downloaded genome, GCA 013398335, was listed as *Nothoprocta ornata* on NCBI but after including newly sequenced material, this *N. ornata* clustered with *N. pentlandii* samples from Peru. After examining the voucher specimen (CORBIDI 168605), the sample was indeed confirmed as *N. pentlandii oustaleti* and not *N. ornata* as indicated in NCBI.

*Assessment of gene tree estimation error*

To investigate the impacts of gene tree estimation error, we first measured gene tree discordance for each dataset using RF distances between gene and species trees. Although gene tree discordance is expected due to the stochastic nature of the coalescent process, we assumed that increases in the mean and variance of RF distances between datasets were attributable to increased gene tree estimation error. In line with this assumption, the UCE100 dataset had the highest RF distances between gene and species trees, whereas the UCE1000 dataset had the lowest mean and variance in RF distances (Figure S22). The BUSCO and UCE300 datasets had intermediate RF distances, with similar means and variances. Interestingly RF distances from the BUSCO dataset were highly variable, with estimated gene trees from some loci showing relatively low (akin to the UCE1000 dataset) and some showing exceptionally high (akin to the UCE100 and UCE300 datasets) RF distances between gene and species trees. A Kruskal-Wallis test confirmed that RF-distances differed significantly among datasets ($X^2$ =3830.5, *df* = 3, $P$ < 0.00001), and a post-hoc Dunn test also revealed that differences between pairwise comparisons of these datasets were significantly different ($P$ < 0.01) except between BUSCO and UCE300 datasets ($P$=0.12). Finally, UCEs mapping to the Z-chromosome (UCE1000) had lower gene tree discordance (mean RF distance between gene and species tree = 32.23.68 ± 9.67) than those on autosomes (mean=38.58 ± 9.67), and this difference was significant ($X^2$=26.65, df=1, P<0.0001), suggesting lower rates of error, ILS and/or introgression on the Z-chromosome. However, there was no difference in RF distances between Z-chromosome UCEs and autosomal UCEs for UCE300 ($X^2$=0.37, df=1, P=0.55) and UCE100 ($X^2$=0.02, df=1, P=0.89) datasets.

We recovered significant differences in the number and proportion of PIS per gene alignment as well as levels of gene tree estimation error among the four datasets. As expected, the UCE100 dataset had the fewest PIS, averaging just 41.10 ± 23.54 PIS per alignment, the UCE300 dataset averaged an intermediate number with a mean of 165.00 ± 62.25 PIS per alignment, and the UCE1000 dataset contained the most PIS with a mean of 682.53 ± 164.17 per alignment. The BUSCO alignments also contained a high number of PIS, although with very high variance, indicating a range of informative and uninformative loci (mean = 529.58 ± 393.29 PIS per locus) (Figure S22). Kruskal-Wallis tests confirmed that differences in both the number ($X^2$ = 3510.1, *df* = 3, $P$ < 0.00001) and proportion ($X^2$ = 6222.1, *df* = 3, $P$ < 0.00001) of PIS significantly differed among datasets. Dunn tests also indicated that differences between pairwise comparisons of these datasets were all significantly different ($P$ adjusted < 0.00001) except that the proportion of PIS per locus did not differ between BUSCO and UCE100 datasets (P=0.38). Within-locus variance in GC content also differed among datasets (Kruskal-Wallis $X^2$ = 229.01, *df* = 3, $P$ < 0.00001), with BUSCOs showing the highest mean and standard deviation (mean = 0.0002 ± 0.0003). A Dunn test confirmed that all pairwise comparisons of datasets differed in GC content variation (*P* adjusted < 0.0001 for all).

To quantify the relationship between alignment information content and gene tree discordance, we modeled RF distances as a function of the number and proportion of PIS per locus using generalized linear models (Figure S23). These models were overall consistent with an expected pattern of a negative association between PIS and RF distance, with lower RF distances in gene trees built from more informative alignments. However, there was variation in the slope and tightness of fit of these models among datasets. For example, the UCE100 and UCE300 datasets had a tight negative association between RF distance and both the number of PIS and percentage of PIS per locus, indicating a large proportion of the variance in RF distance can be explained by alignment information content. The same negative association was revealed for UCE1000 dataset, but this relationship was noisier with a lower $R^2$, indicating a weaker association. These differences were better visualized when the three UCE datasets were combined, revealing a strong and tightly fitting negative logarithmic relationship between the number PIS and RF distance ($R^2$=0.92). We recovered a similar relationship for the BUSCO dataset using the number of PIS as the predictor variable, but this was noisier than UCEs ($R^2$=0.50), especially when using the proportion of PIS as the predictor variable ($R^2$=0.01). This result was robust, holding true for BUSCO datasets that were filtered to resemble UCE-like datasets and after filtering outliers in GC content variation (Figure S24).

*Breakpoint analysis to find the optimal number of reticulate nodes*

The slope of the segmented model was significant and tightly fitting for both autosomal UCEs (Breakpoint = 3, Adj. $R^2$ = 0.97, P < 0.0001) and Z-linked UCEs (Breakpoint = 1, Adj. $R^2$ = 0.98, P < 0.0001).

### Discussion

*Comparisons of estimation error between data types and the benefits of whole genome phylogenomics*

We measured rates of gene tree discordance using RF distances between gene trees and the inferred species tree with the assumption that, if RF distance was strongly associated with alignment information content, then much of the variation in RF distance is driven by stochastic error, rather than true variance in gene histories. Although the $R^2$ was small for the UCE1000 dataset, the overall effect was most evident when the three UCE datasets were combined into a single analysis, which showed asymptotic convergence on relatively low mean and variance RF distances after reaching about 500 PIS (Figure S23). The asymptotic shape of this relationship implies that as PIS are added to an alignment, the resulting gene trees converge on a level of discordance that may be approaching real biological signal (i.e., ILS and/or introgression) rather than methodological or data-driven artifacts. For example, 92% of the variance in RF distances in the combined UCE regression (Figure 23 top right) is explained by a PIS, but for the UCE1000 dataset, only 29% of the variance in RF is explained by PIS. Given the asymptotic shape of the curve (log-linear), we would argue that relatively long UCE's have reduced stochastic error relative to the other datasets. However, it is admittedly quite difficult to separate this hypothesis from one where analyzing longer sequences (that include more PIS) may lead to the inclusion of multiple c-genes, which could then drive convergence in gene tree topologies by mimicking the effects of concatenation. Although some level of gene tree discordance is expected in *all* phylogenomic datasets, even if one assumes the impossible, where empirical gene tree estimation error is absent (Maddison 1997; Gatesy and Springer 2014; Edwards et al. 2016), it is reasonable to assume that evaluating differences in the mean and variance in RF distances among datasets can help identify differences in gene tree estimation error given the results of the generalized linear models. Nevertheless, data biases and stochastic error had relatively little impact on most of our phylogenetic results; only the

MSC results were impacted by stochastic error; UCE100 and UCE300 MSC trees were quite different from other topologies.

We scrutinized orthologous data harvested from whole genome assemblies to identify the characteristics of phylogenomic datasets that might lead to lower bias. We found that, as expected, short alignments with relatively few PIS had increased RF distances between gene tree and species trees, a finding consistent with expectations of stochastic error described in other studies (Meiklejohn et al. 2016; Burbrink et al. 2020). We therefore used the UCE1000 dataset for analysis of ILS and introgression to avoid confounding effects of putatively erroneous gene trees in the other datasets. Although it is likely that some effects of stochastic error remain (e.g., $R^2 = 0.29$ in the UCE1000 log-linear model), we argue that much of the gene tree discordance found in the UCE1000 dataset is likely representative of biological processes like ILS and introgression. Although all datasets conformed to the expected negative association between PIS and gene tree estimation error (Figure S23), the association was somewhat noisier for BUSCOs than for UCEs pointing toward potential additional sources of discordance.

Some studies have found that coding sequences perform poorly relative to putatively neutral loci due to potential model misspecification, poor absolute model fit, or skewed GC content variation (Reddy et al. 2017; Stiller et al. 2024). We tested whether some reduction in $R^2$ relative to UCEs could be associated with such biases, but after filtering the BUSCO dataset for genes with GC content variation and other characteristics comparable to UCEs, we did not see an increase in $R^2$. Because BUSCOs are constructed from spliced exons that may be separated from each other by introns of tens to hundreds-of-thousands of base pairs in length, it is also possible that the increased rate of gene tree error is driven by the inclusion of many freely recombining *c*-genes in each gene alignment, which is known to impact phylogenetic estimation (Doyle 1995, 1997; Schierup and Hein 2000; Hobolth et al. 2007). This phenomenon could be unlikely in this case because increasing the number of *c*-genes used to build each gene tree should cause gene tree topologies to converge on the concatenated topology. Although we found concordance between concatenated and MSC BUSCO trees, the high variance in RF distances between BUSCO gene and species trees, is not consistent with this expectation.

Although we derived datasets from whole genome sequences, the UCE100 and UCE300 datasets are likely to closely match datasets obtained from typical target capture approaches (Smith et al. 2014; McCormack et al. 2016; Musher and Cracraft 2018; Tea et al. 2021) and may therefore be indicative of rates of stochastic error in datasets derived from those protocols. For example, many studies utilize sequences from degraded DNA such as historical museum specimens, which may result in many relatively short UCE alignments (McCormack et al. 2016), but even UCE-enriched datasets generated from fresh tissues typically only capture about 300 bp of flanking region around the conserved UCE core.

Given these findings, common phylogenomic approaches that sequence loci with intermediate or low levels of information content such as sequence capture of UCEs may be insufficient to resolve the relationships of taxa that evolved rapidly, such as adaptive radiations, where rates of ILS due to rapid diversification are expected to be intense (Gatesy and Springer 2014; Mclean et al. 2019). Even in this study, which utilized multiple data types and loci of varying lengths, the phylogenetic placement of one rapidly diverged species, *C. atrocapillus,* was equivocal. In these situations, the number of variable sites may still be too few to resolve gene trees with short branches, even with whole genome data.

### Supplemental Tables and Figures:

**Table S1: A list of samples used in this study including statistics for sequence coverage and genome completeness (supplemental file only, not shown)**

| Fossil | Stratigraphic level | Estimated age (Ma) | Phylogenetic position justification | Prior Details | Citations |
|---|---|---|---|---|---|
| *Diogenornis fragillis* | Itaboraian-Casamayoran | 53–48.6<br><br>Initial tip date: NA | Multiple synapomorphies with Rheidae place this fossil in Rheiformes | Lognormal MRCA constraint on age of "Higher ratites" Offset=55, M=7, S=1 | Alvarenga 1983, Almeida et al. 2022 |
| Undescribed fossil Macn-sc-3610 | Monte observacion, Santa Cruz Formation | 18–15.2<br><br>Initial tip date: 16.6 | Phylogenetic analysis of morphological data placed this fossil as sister to crown *Crypturellus* | Constrained monophyly of *Crypturellus* + stem fossils, thus allowing this fossil to freely vary within stem and crown *Crypturellus* | Bertelli and Chiappe 2005, Bertelli et al. 2014, Almeida et al. 2022 |
| Undescribed fossil Macn-sc-3613 | Monte observacion, Santa Cruz Formation | 18–15.2<br><br>Initial tip date: 16.6 | Phylogenetic analysis of morphological data placed this fossil as sister to crown *Crypturellus* | Constrained monophyly of *Crypturellus* + stem fossils, thus allowing this fossil to freely vary within stem and crown *Crypturellus* | Bertelli and Chiappe 2005, Bertelli et al. 2014, Almeida et al. 2022 |
| *Crypturellus reai* | Cañadón de las Vacas, Santa Cruz Formation | 17.5–16.3<br><br>Initial tip date: 16.9 | Previous phylogenetic analysis was equivocal, placing this specimen either in stem or crown *Crypturellus* | Constrained monophyly of *Crypturellus* + stem fossils, thus allowing this fossil to freely vary within stem and crown *Crypturellus* | Bertelli et al. 2014, Almeida et al. 2022 |
| *"Eudromia"* sp. | Cerro Azul Formation at Salinas Grandes | 7.2–6<br><br>Initial tip date: 6.6 | Previous phylogenetic analysis placed this fossil as sister to *Eudromia* + *Tinamotis* | Constrained phylogenetic position of "*Eudromia* sp." thus constraining the position sister to *Tinamotis*+*Eudromia* | Bertelli et al. 2014, Almeida et al. 2022 |
| *Eudromia olsoni* | Farola Monte Hermoso Montehermosan | 5–4.5<br><br>Initial tip date: 4.75 | Previous phylogenetic analysis placed this fossil in the genus *Eudromia* | Constrained monophyly of extant *Eudromia* + fossil allowing the fossil to freely vary within crown *Eudromia* | Bertelli et al. 2014, Almeida et al. 2022 |

| | | | | | |
|---|---|---|---|---|---|
| *Nothura parvula* | Chapadmalal Formation | 5–4.5<br><br>Initial tip date: 4.75 | Previous phylogenetic analysis was equivocal, placing this species either in *Nothura* or as sister to *Nothura* + *Nothoprocta* + *Rhynchotus*.<br>Lack of synapomorphies with *Nothoprocta* + *Rhynchotus* suggest it is a stem or crown *Nothura.* | Constrained monophyly of *Nothoprocta* + *Rhynchotus*, thus allowing this fossil to freely vary within stem and crown *Nothura*, and stem *Nothoprocta* + *Rhynchotus* | Bertelli et al. 2014; Almeida et al. 2022 |

**Table S2:** A list of all tip fossils used in the Beast analysis along with justification for their constrained positions, prior distributions and citations.

| **Parameter Name** | **Prior** | **Details** |
|---|---|---|
| ORCRates | Log Normal | Initial=0.5, M=1.0, S=0.2, Offset=0.0 |
| ORCSigma | Exponential | Initial=0.2, M=0.3337,Offset=0.0 |
| ORCucldMean | Exponential | Initial=0.001, M=10.0, Offset=0.0 |
| rhoFBD | Uniform | Initial=0.75, Lower=0.5, Upper=1.0, Offset=0.0 |
| SamplingProportionFBD | Exponential | Initial=0.25, M=0.25, Offset=0.0 |
| MRCA | No prior | Constrained monophyly for 14 clades to improve accuracy of the phylogenetic results given the relatively small number of loci used. These are marked with white circles in Figure 1 of the main text. |

**Table S3:** Non-default priors used in the beast analysis

| Dataset | Number of loci | Total concatenated length |
|---|---|---|
| UCE100 complete | 2,712 | 879,985 |
| UCE100 75% complete | 4,631 | 1,482,732 |
| UCE300 complete | 2,702 | 1,864,316 |
| UCE300 75% complete | 4,621 | 3,144,625 |
| UCE1000 complete | 2,628 | 4,643,253 |
| UCE1000 75% complete | 4,514 | 7,846,363 |
| UCE1000 autosomes 75% complete | 4,244 | 7,384,066 |
| UCE1000 Z-chromosome 75% complete | 304 | 677,088 |
| BUSCOs complete | 2,274 | 3,103,102 |
| BUSCOs 75% complete | 7,414 | 10,083,122 |

**Table S4:** Number of alignments and total concatenated length for each dataset.

| Dataset | Model | Adj. R-squared | slope | P | AIC |
|---|---|---|---|---|---|
| BUSCOs | linear (#PIS) | 0.3387 | -0.0305809 | <0.0001 | 19291 |
| | logarithmic (#PIS) | 0.497 | -20.6024 | <0.0001 | 18669 |
| | linear (%PIS) | 0.001863 | -7.36 | 0.0396 | 20228 |
| | logarithmic (%PIS) | 0.01044 | -6.684 | <0.0001 | 20208 |
| UCE100Flank | linear (#PIS) | 0.7797 | -0.759178 | <0.0001 | 19897 |
| | logarithmic (#PIS) | 0.6933 | -25.8442 | <0.0001 | 20794 |
| | linear (%PIS) | 0.7747 | -237.6429 | <0.0001 | 19959 |
| | logarithmic (%PIS) | 0.6933 | -25.8442 | <0.0001 | 20843 |
| UCE300Flank | linear (#PIS) | 0.636 | -0.254330 | <0.0001 | 21092 |
| | logarithmic (#PIS) | 0.7253 | -37.7134 | <0.0001 | 20331 |
| | linear (%PIS) | 0.6039 | -160.9276 | <0.0001 | 21320 |
| | logarithmic (%PIS) | 0.705 | -35.7484 | <0.0001 | 20523 |
| UCE1000Flank | linear (#PIS) | 0.24 | -0.028939 | <0.0001 | 18683 |
| | logarithmic (#PIS) | 0.2895 | -18.5317 | <0.0001 | 18506 |
| | linear (%PIS) | 0.1312 | -39.8117 | <0.0001 | 19035 |
| | logarithmic (%PIS) | 0.2287 | -9.0358 | <0.0001 | 18927 |
| UCE Combined | linear (#PIS) | 0.6814 | -0.108918 | <0.0001 | 24575 |
| | logarithmic (#PIS) | 0.9197 | -28.6239 | <0.0001 | 20828 |
| | linear (%PIS) | 0.7765 | -248.5049 | <0.0001 | 23611 |
| | logarithmic (%PIS) | 0.791 | -48.0636 | <0.0001 | 23428 |

**Table S5:** Model tests of log versus linear fits for all generalized linear models. This model test was done twice for each dataset, once using the number of parsimony informative sites per locus (#PIS) and once using the percentage of parsimony informative sites per locus (%PIS). Rows highlighted in gray indicate the best model for each paired model test.

**Table S6: Clade ages from this and other studies (supplemental file only, <u>not shown</u>)**

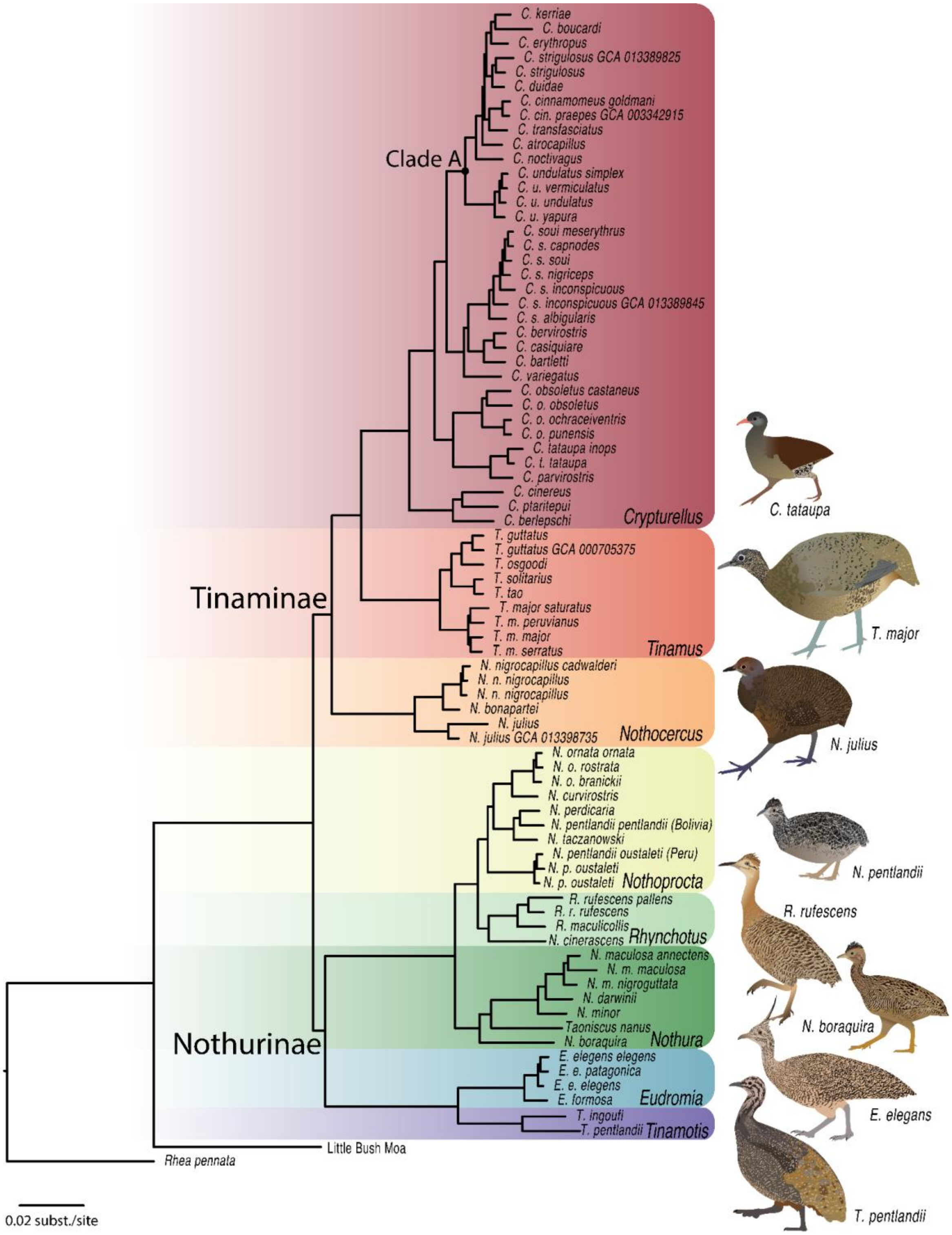

**Figure S1: Concatenated Phylogeny of tinamous based on UCEs with 1,000 bp of flanking sequence.** All nodes have bootstrap value of 100% except where noted. Note relatively short internodal branch lengths in Clade A. Tinamou illustrations by TAC.

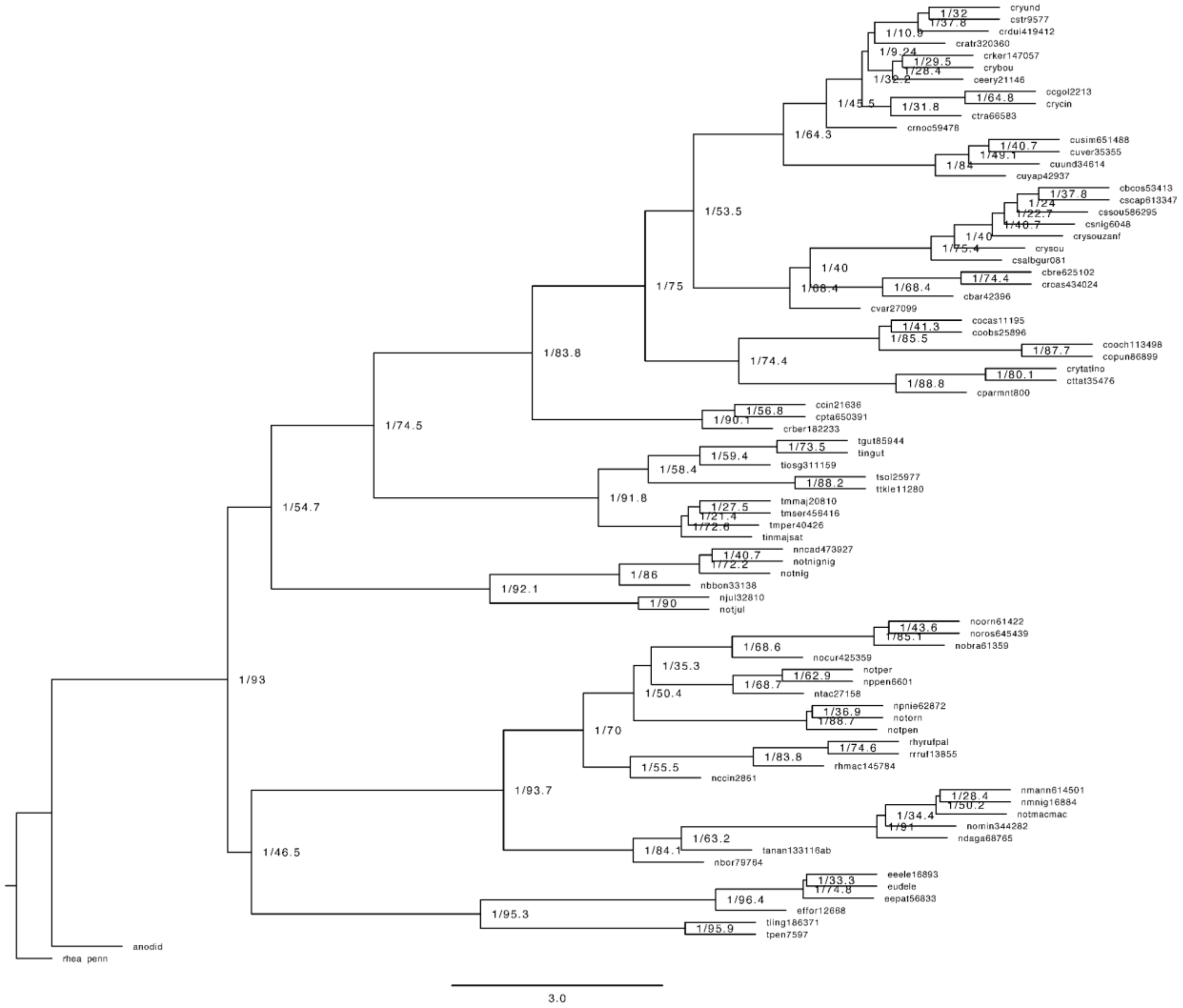


**Figure S2: BUSCOs 75% complete Astral phylogeny.**

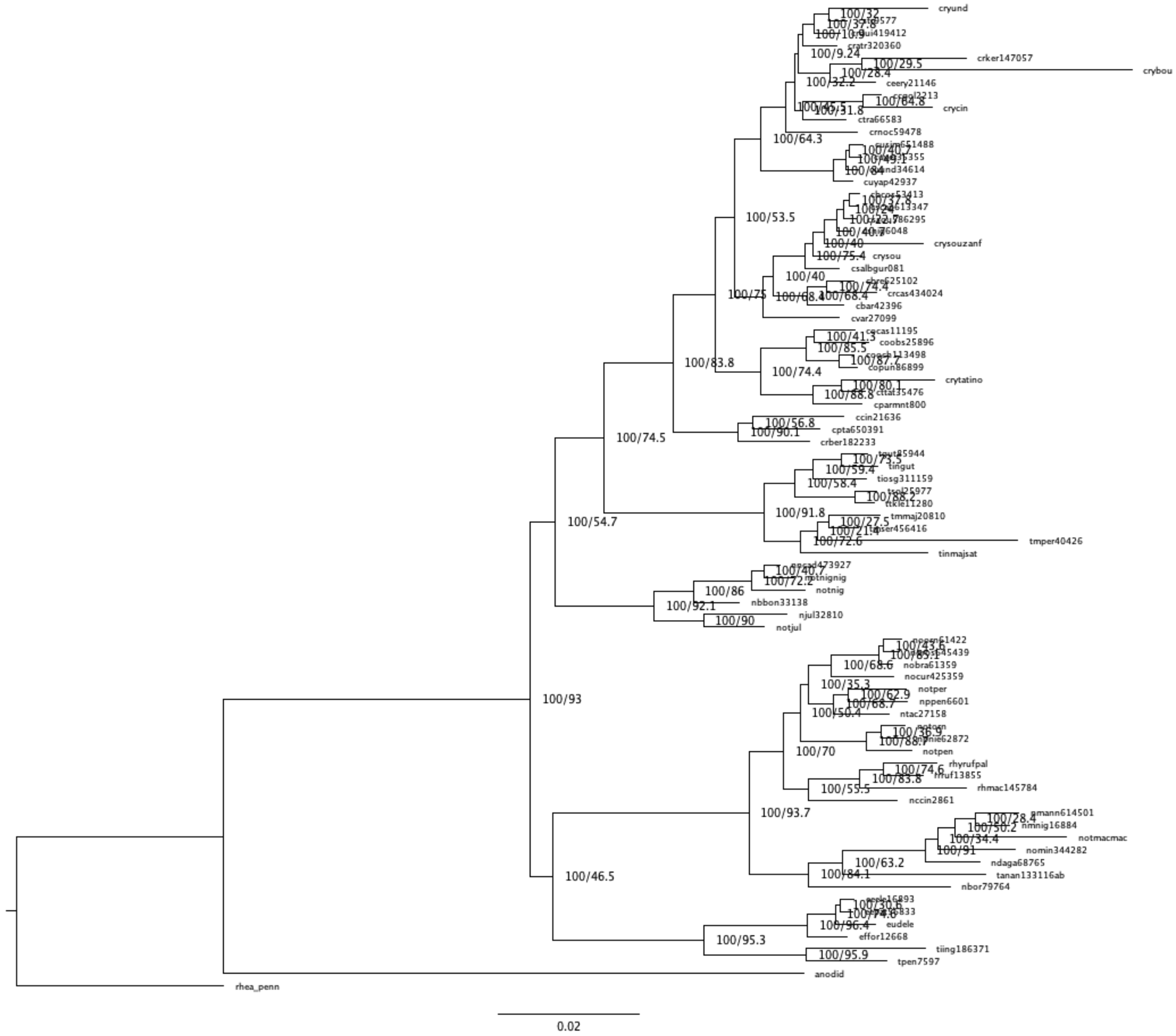

**Figure S3: BUSCOs 75% iqtree phylogeny**

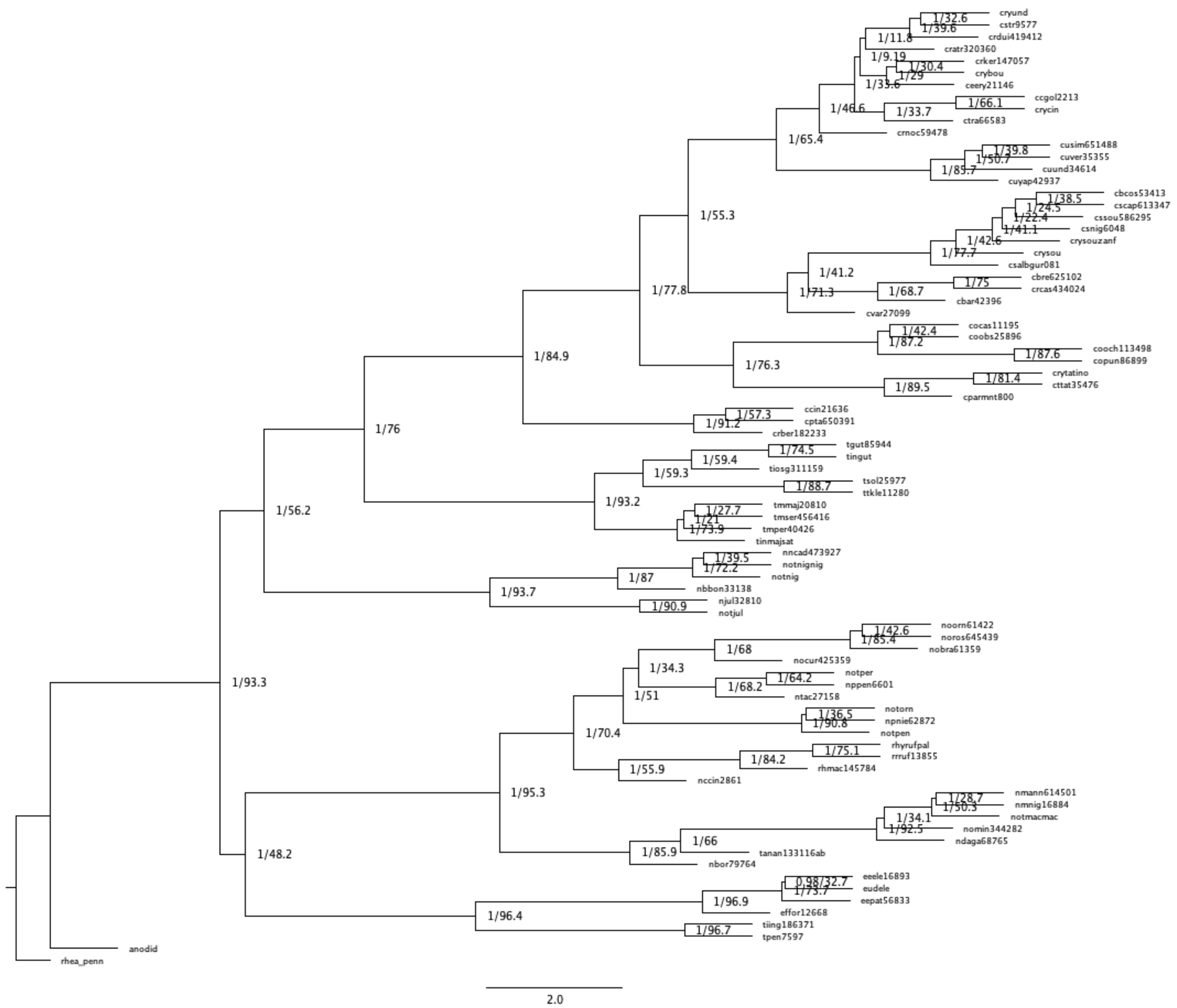


**Figure S4: BUSCOs 100% Astral phylogeny**

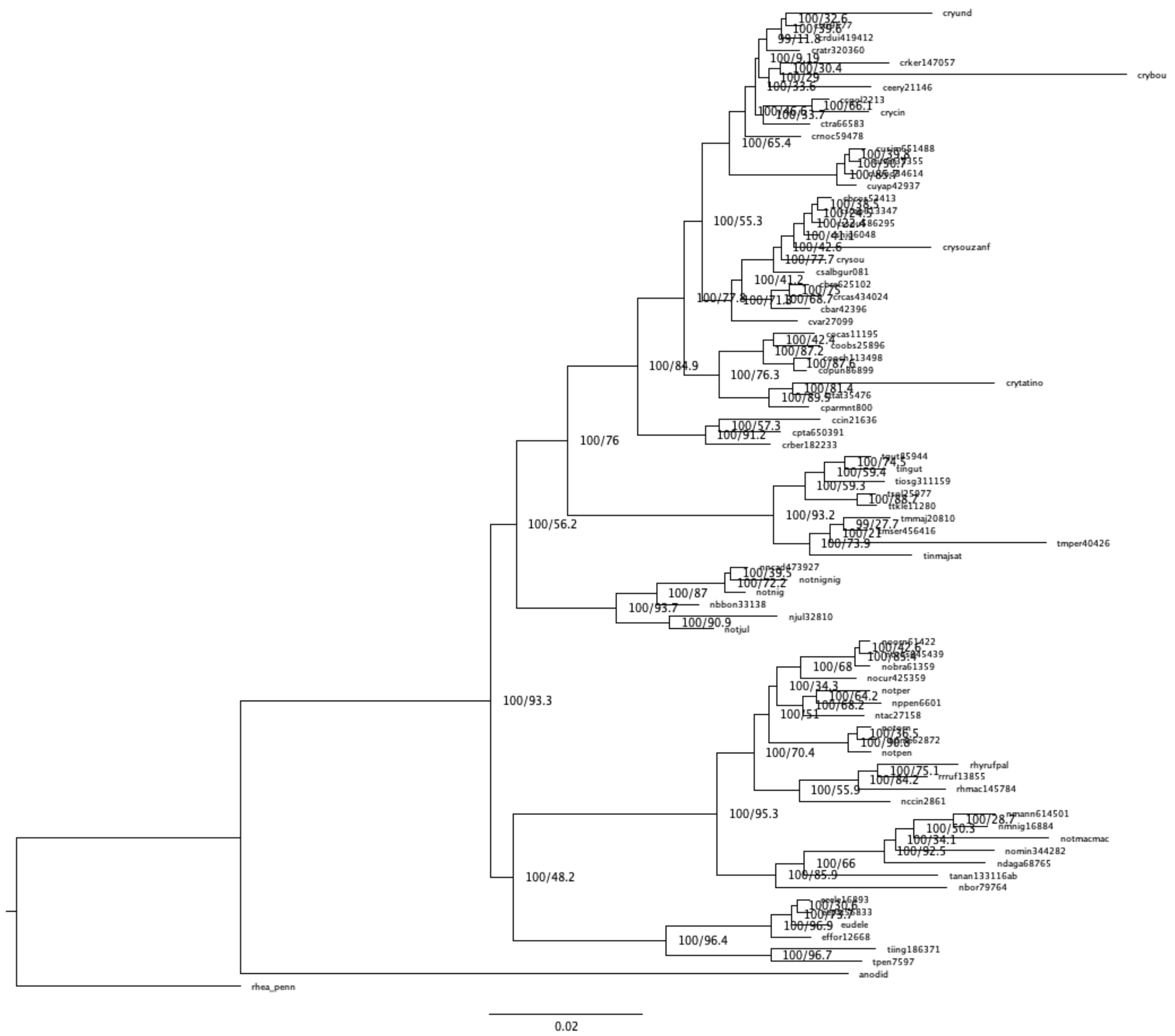


**Figure S5: BUSCOs 100% iqtree phylogeny**

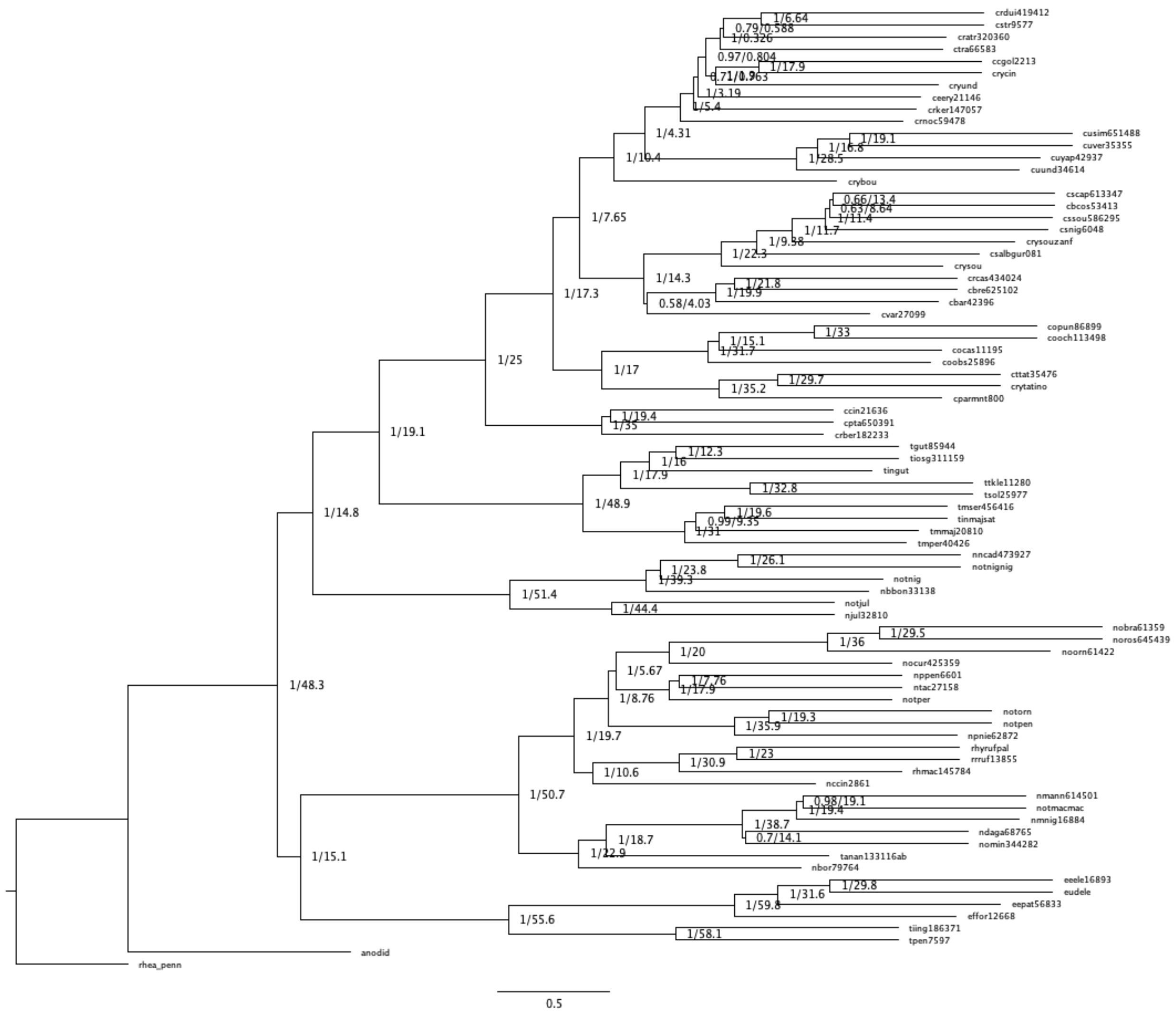

**Figure S6: UCE100 75% Astral phylogeny**

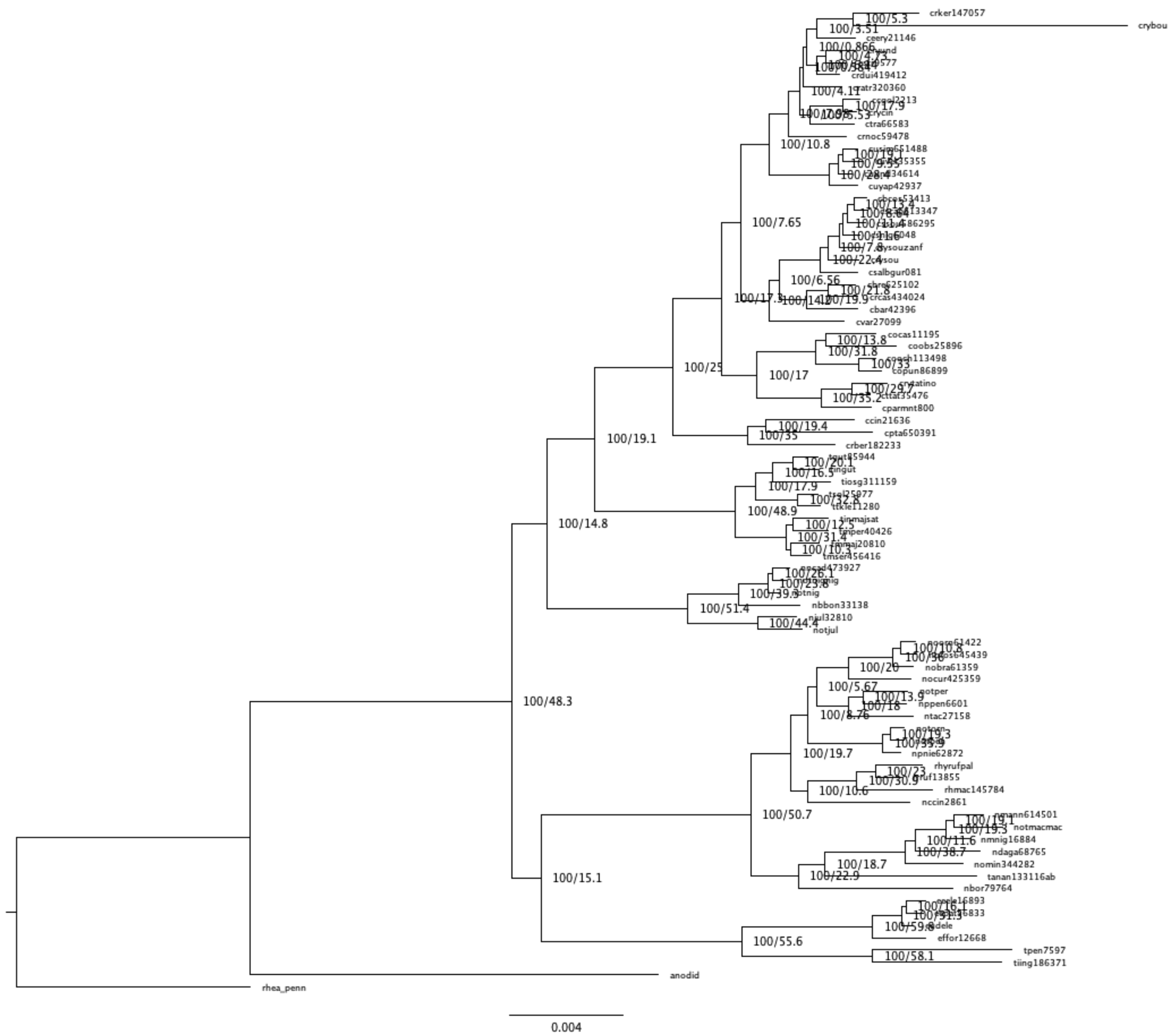

**Figure S7: UCE100 75% iqtree phylogeny**

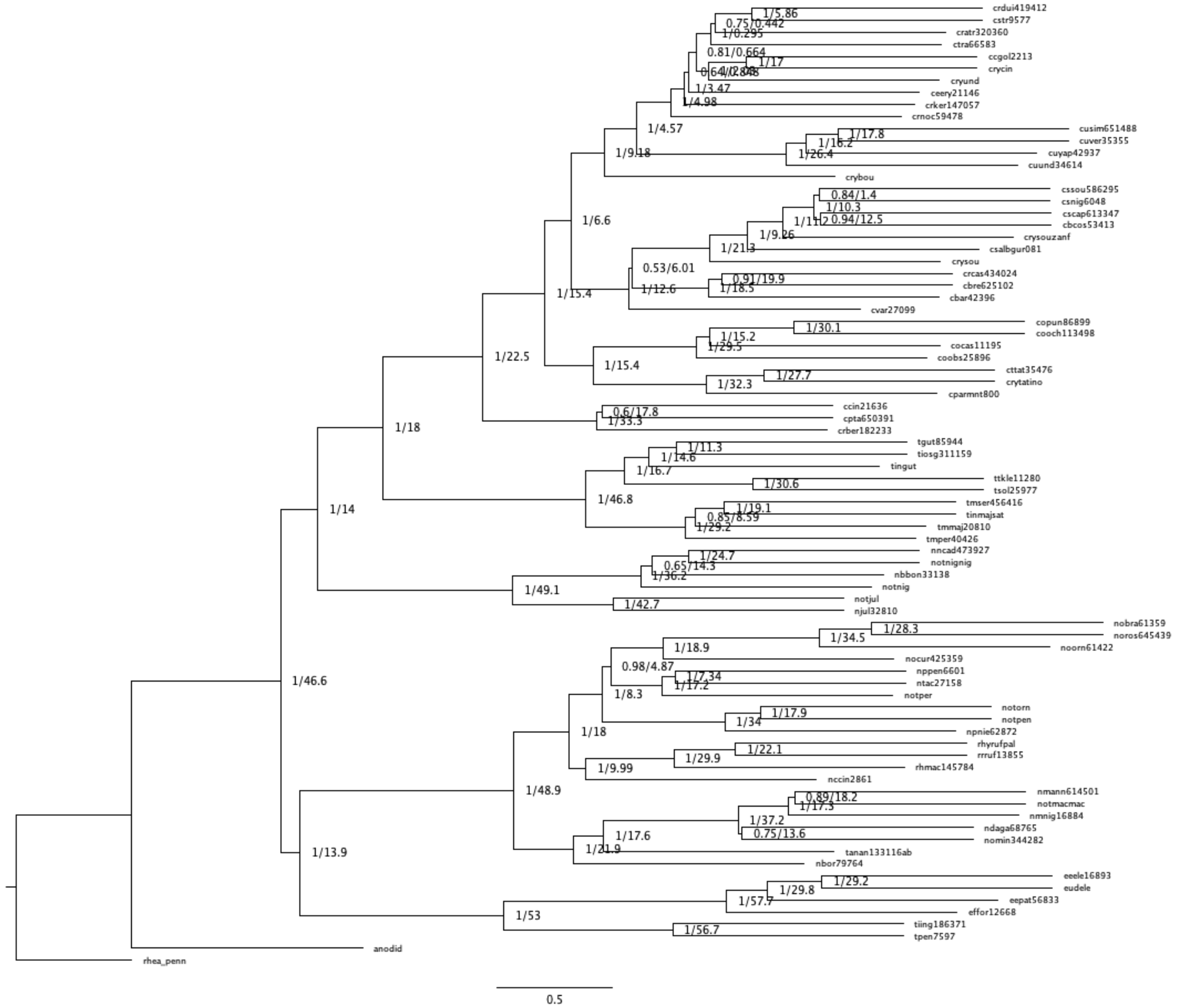

**Figure S8: UCE100 100% Astral phylogeny**

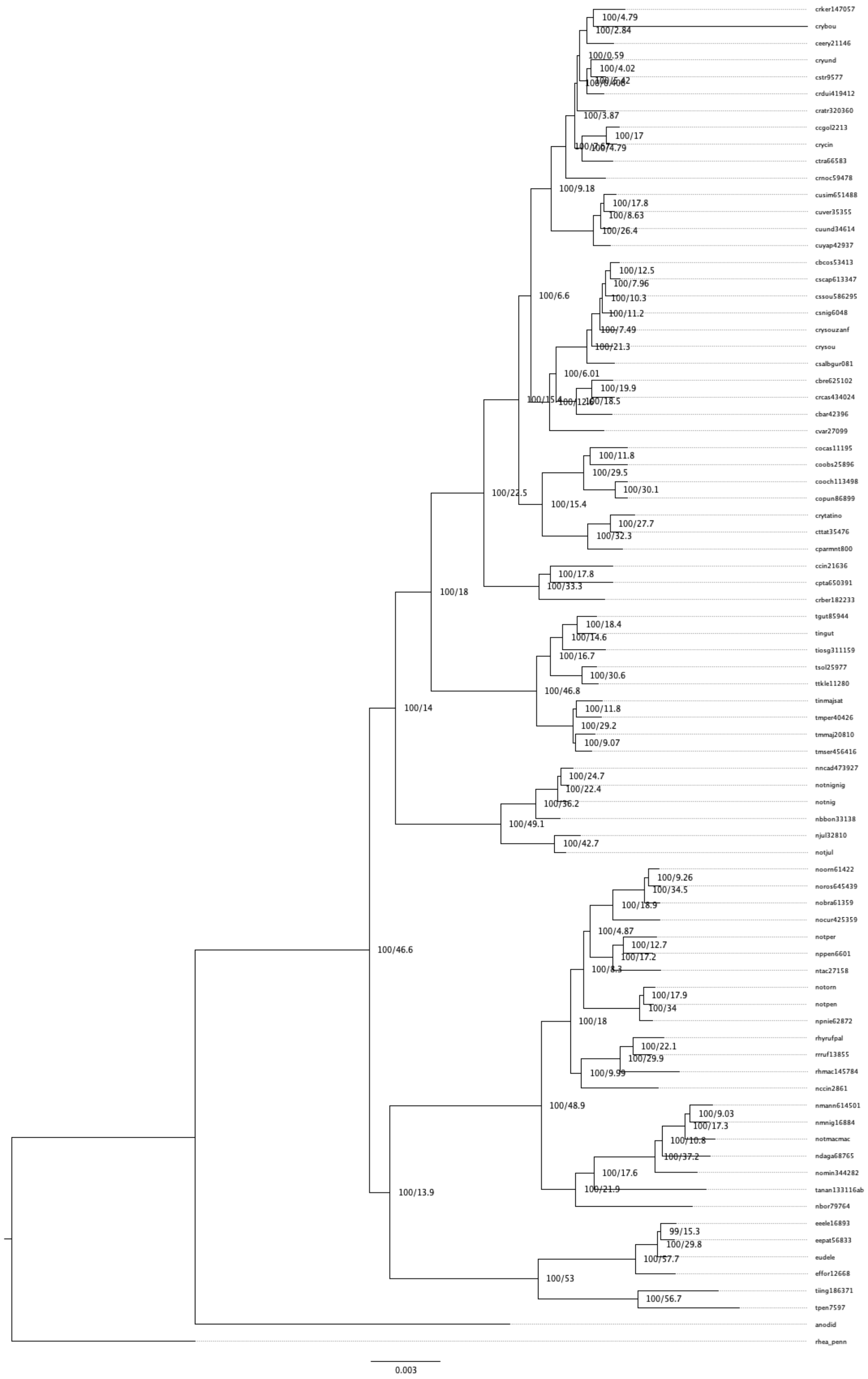
crker147057
crybou
ceery21146
cryund
cstr9577
crdui419412
cratr320360
ccgol2213
crycin
ctra66583
crnoc59478
cusim651488
cuver35355
cuund34614
cuyap42937
cbcos53413
cscap613347
cssou586295
csnig6048
crysouzanf
crysou
csalbgur081
cbre625102
crcas434024
cbar42396
cvar27099
cocas11195
coobs25896
cooch113498
copun86899
crytatino
cttat35476
cparmnt800
ccin21636
cpta650391
crber182233
tgut85944
tingut
tiosg311159
tsol25977
ttkle11280
tinmajsat
tmper40426
tmmaj20810
tmser456416
nncad473927
notnignig
notnig
nbbon33138
njul32810
notjul
noorn61422
noros645439
nobra61359
nocur425359
notper
nppen6601
ntac27158
notorn
notpen
npnie62872
rhyrufpal
rrruf13855
rhmac145784
nccin2861
nmann614501
nmnig16884
notmacmac
ndaga68765
nomin344282
tanan133116ab
nbor79764
eeele16893
eepat56833
eudele
effor12668
tiing186371
tpen7597
anodid
rhea_penn
100/4.79
100/2.84
100/0.59
100/4.02
100/3.87
100/17
100/4.79
100/9.18
100/17.8
100/8.63
100/26.4
100/6.6
100/12.5
100/7.96
100/10.3
100/11.2
100/7.49
100/21.3
100/6.01
100/19.9
100/18.5
100/22.5
100/11.8
100/29.5
100/30.1
100/15.4
100/27.7
100/32.3
100/18
100/17.8
100/33.3
100/18.4
100/14.6
100/16.7
100/30.6
100/46.8
100/11.8
100/29.2
100/9.07
100/14
100/24.7
100/22.4
100/36.2
100/49.1
100/42.7
100/46.6
100/9.26
100/34.5
100/18.9
100/4.87
100/12.7
100/17.2
100/8.3
100/17.9
100/34
100/18
100/22.1
100/29.9
100/9.99
100/48.9
100/9.03
100/17.3
100/10.8
100/37.2
100/17.6
100/21.9
100/13.9
99/15.3
100/29.8
100/57.7
100/53
100/56.7
0.003

### Figure S9: UCE100 100% iqtree phylogeny

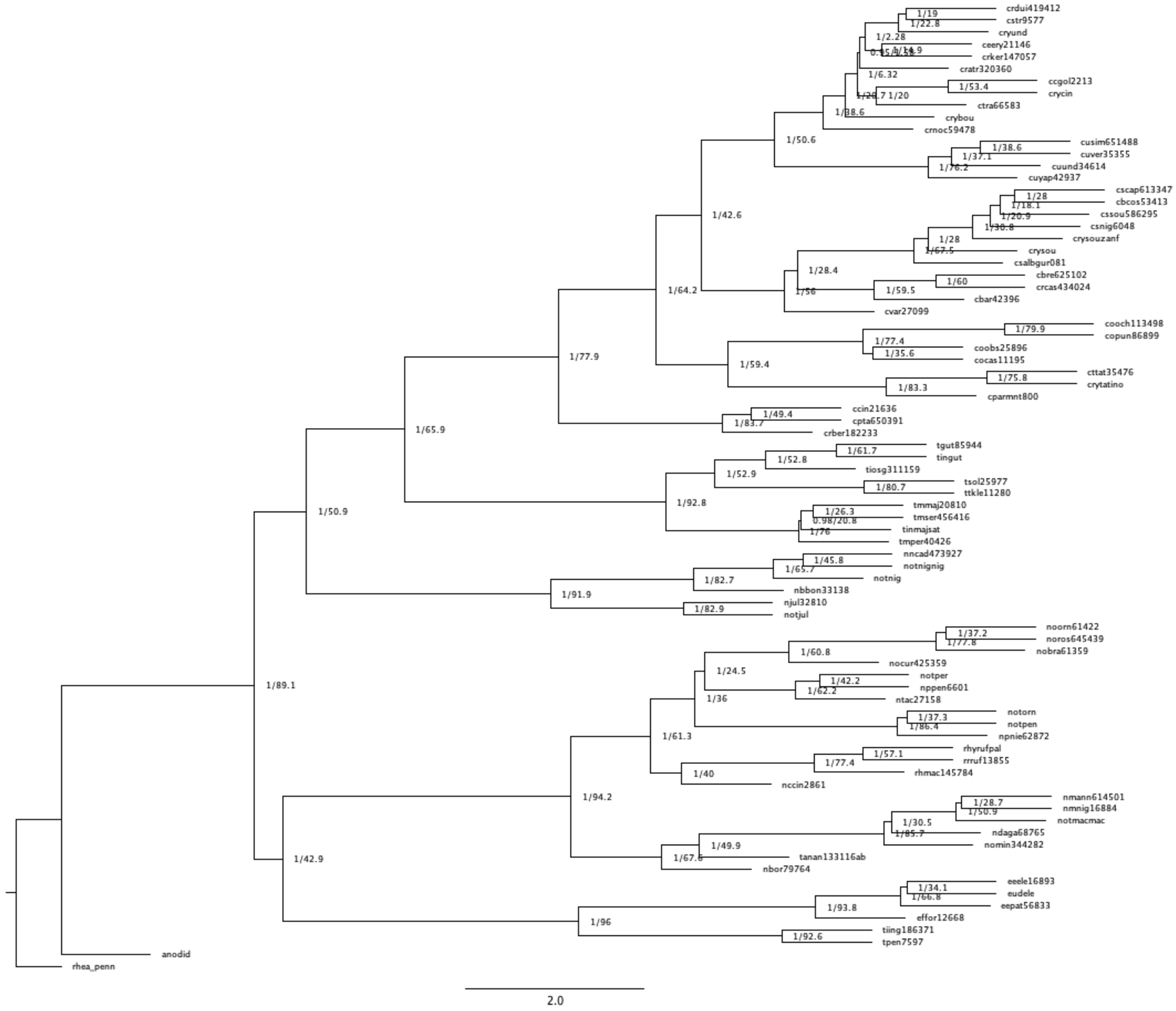


### Figure S10: UCE300 75% ast phylogeny

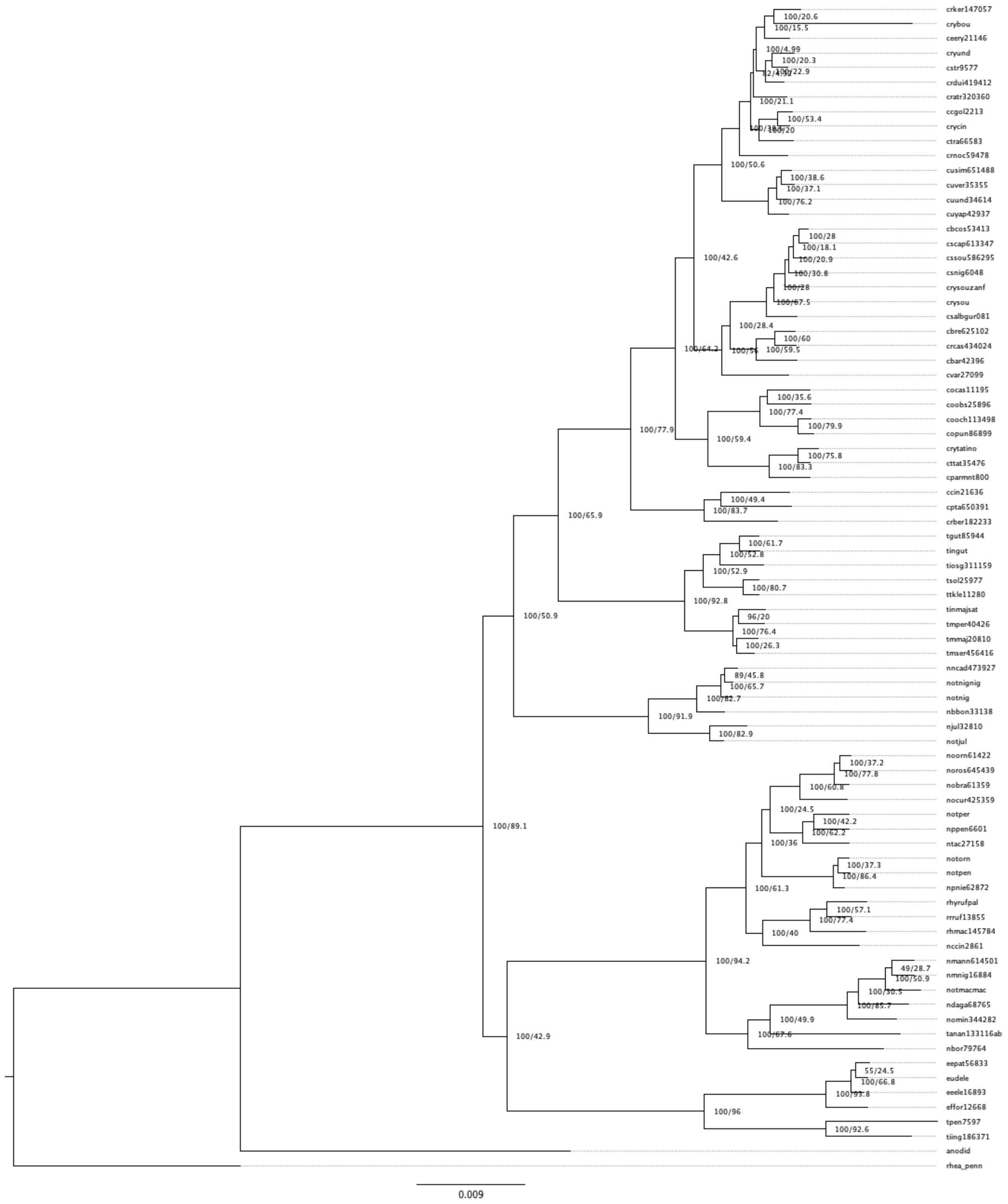


**Figure S11: UCE300 75% iqtree phylogeny**

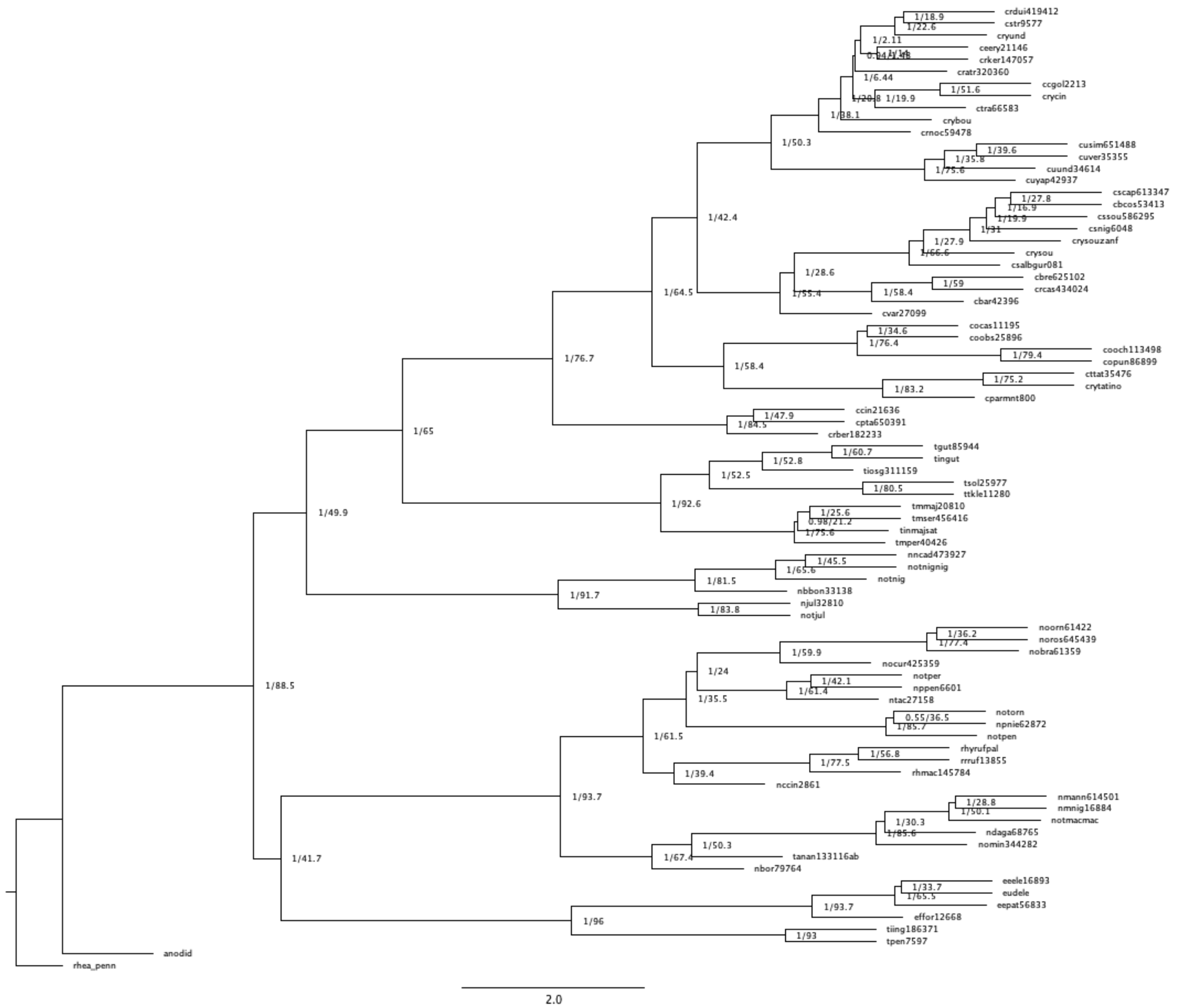


**Figure S12: UCE300 100% Astral phylogeny**

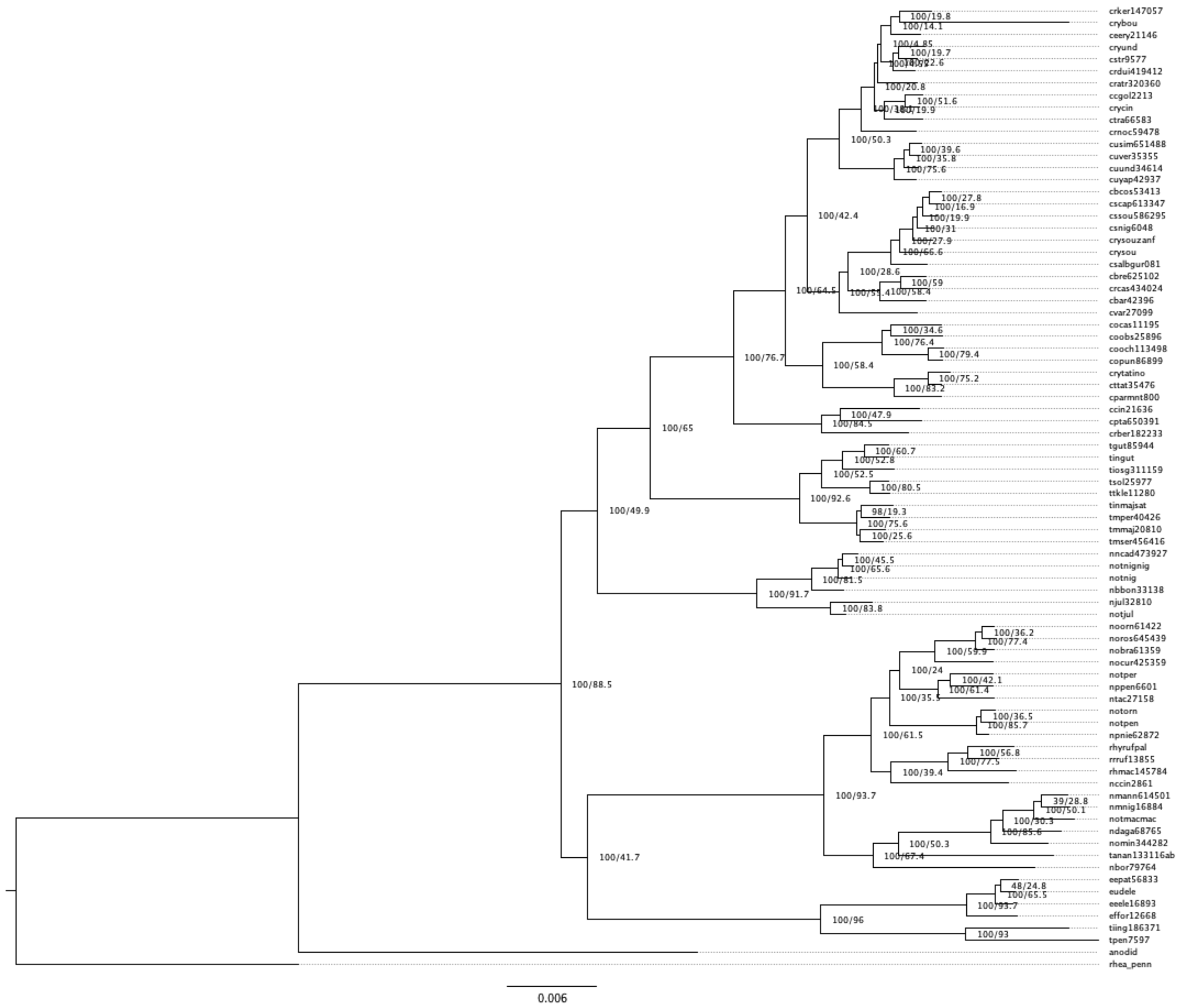


**Figure S13: UCE300 100% iqtree phylogeny**

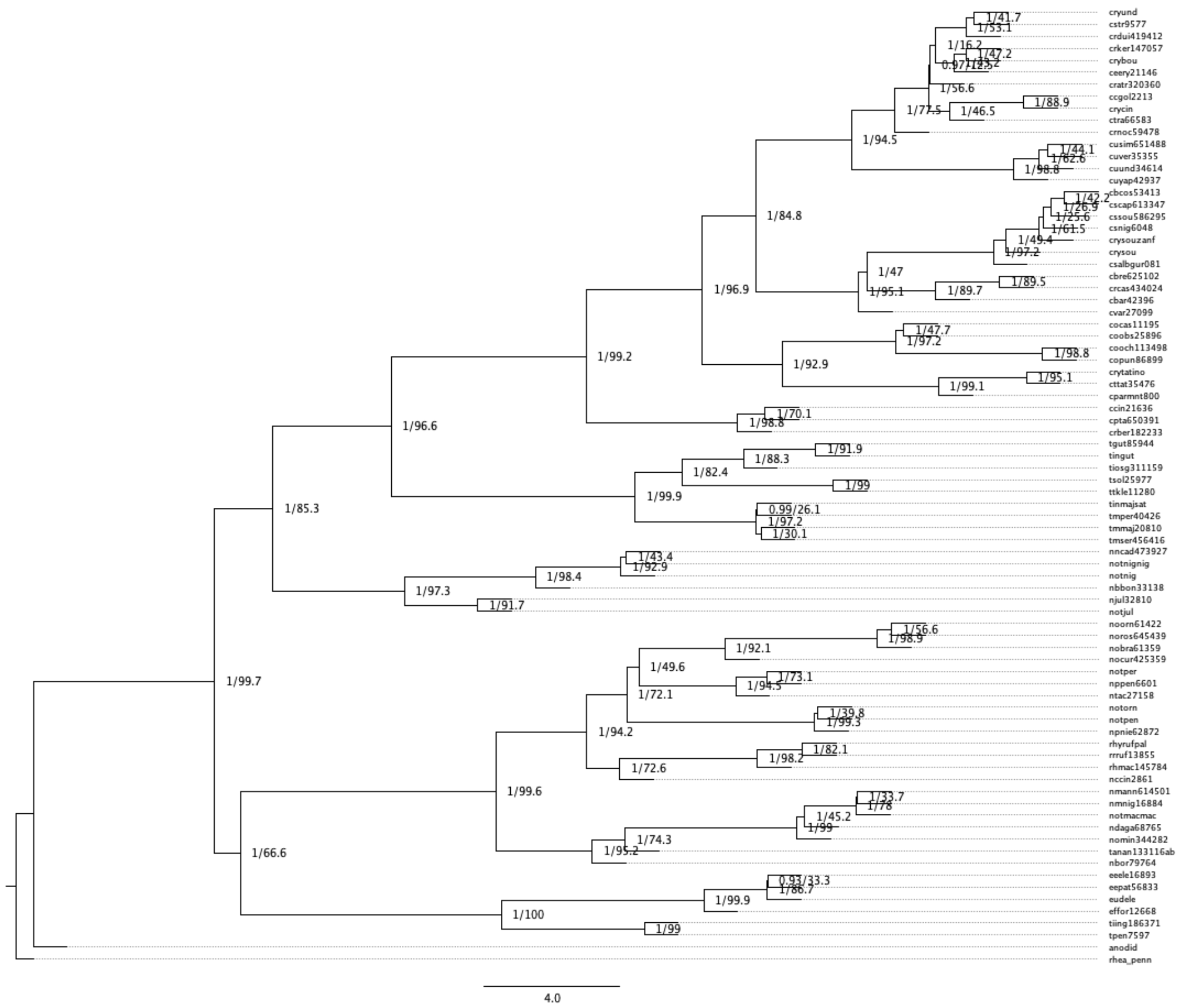

**Figure S14: UCE1000 autosomes 75% Astral phylogeny**

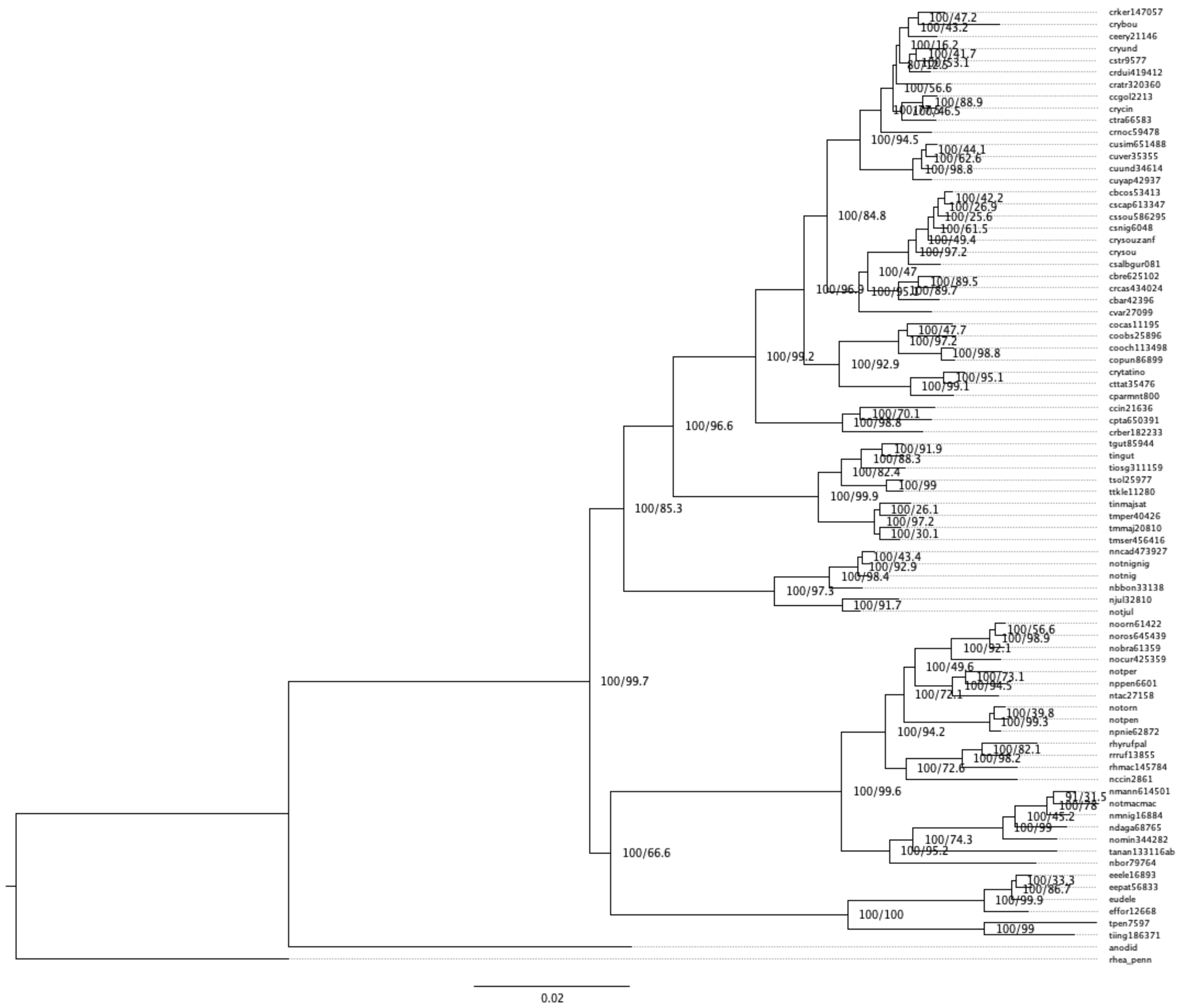


**Figure S15: UCE1000 autosomes 75% iqtree phylogeny**

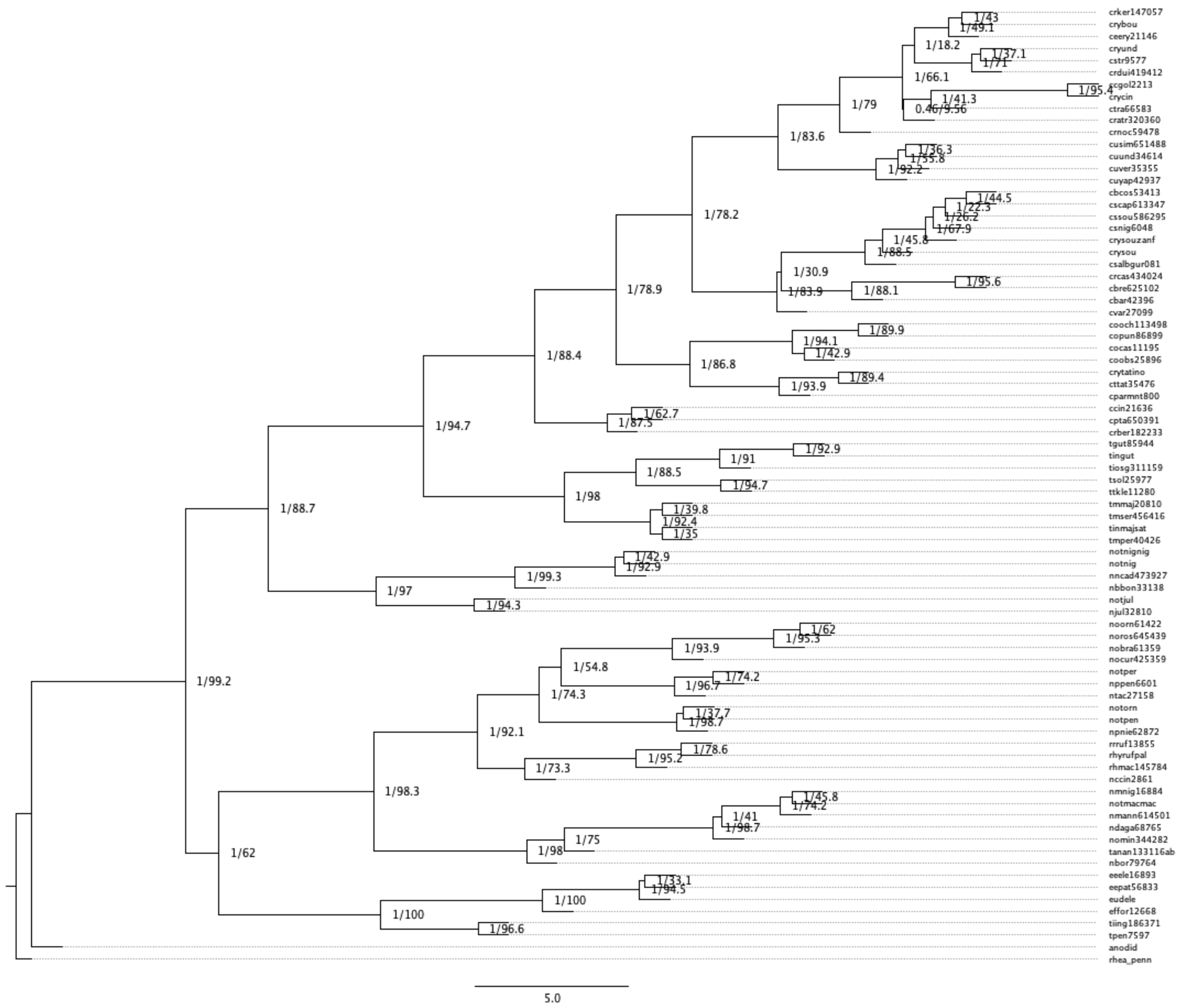


**Figure S16: UCE1000 Z chromosome 75% Astral phylogeny**

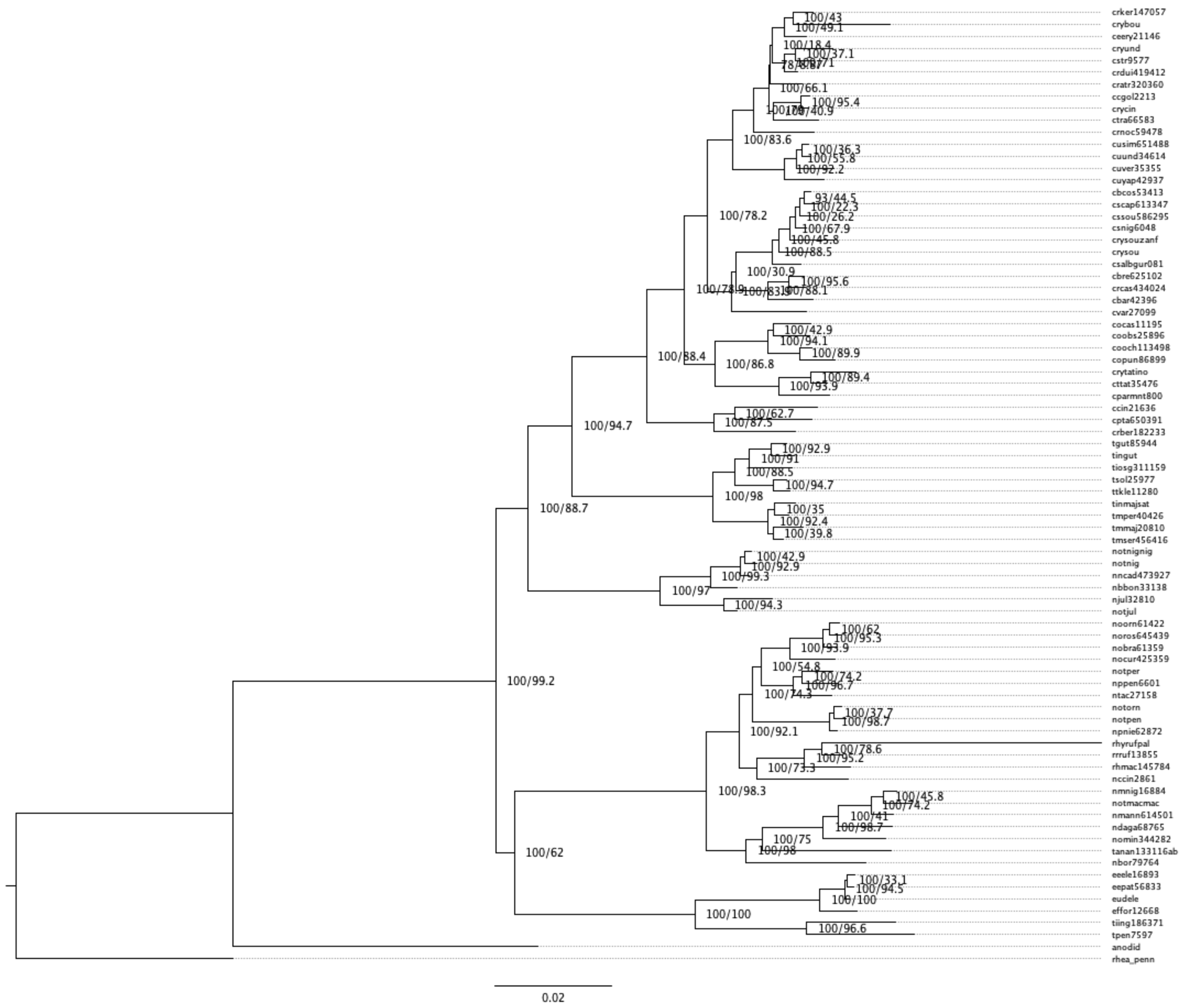

**Figure S17: UCE1000 Z chromosome 75% iqtree phylogeny**

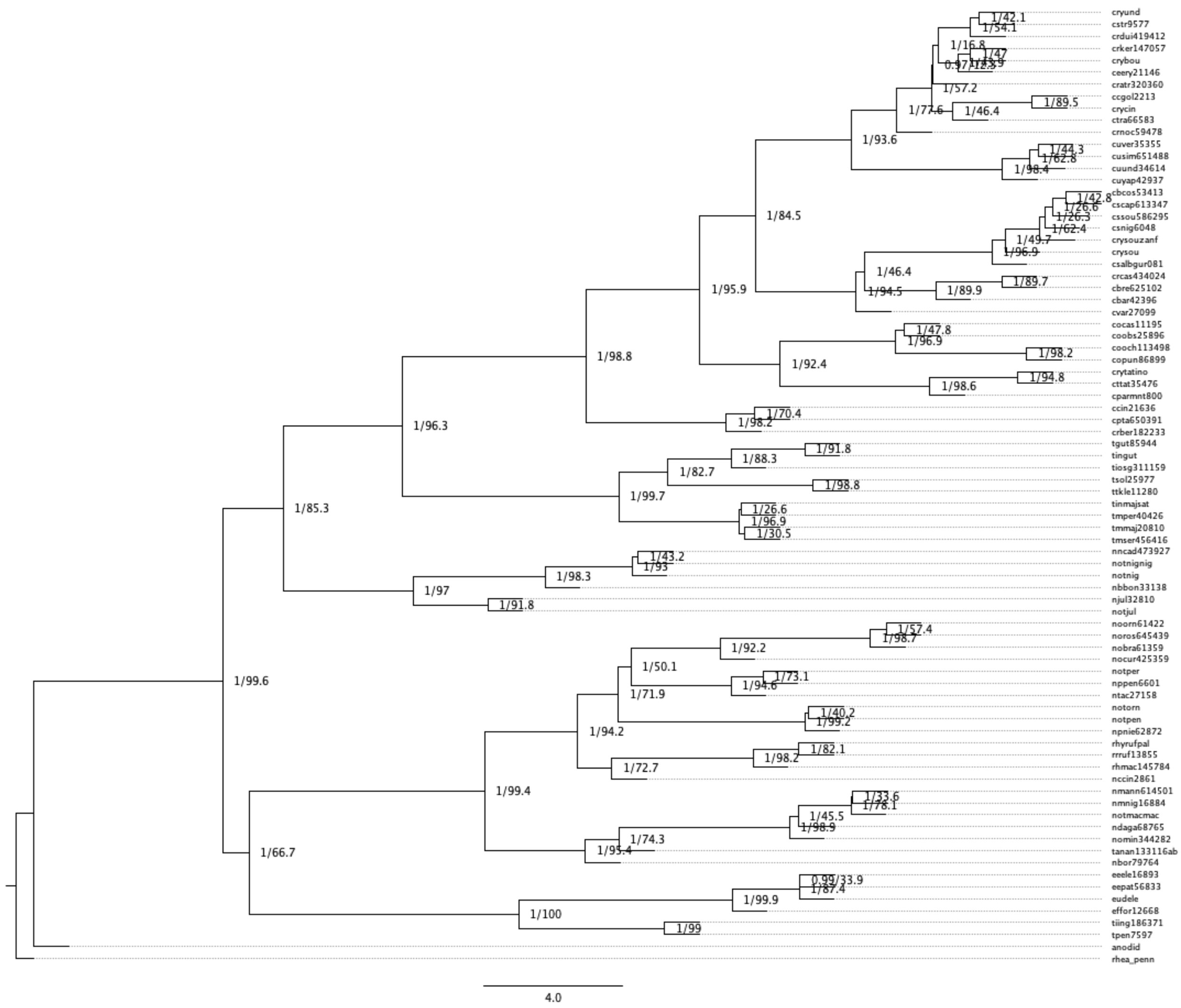

**Figure S18: UCE1000 75% whole dataset Astral phylogeny**

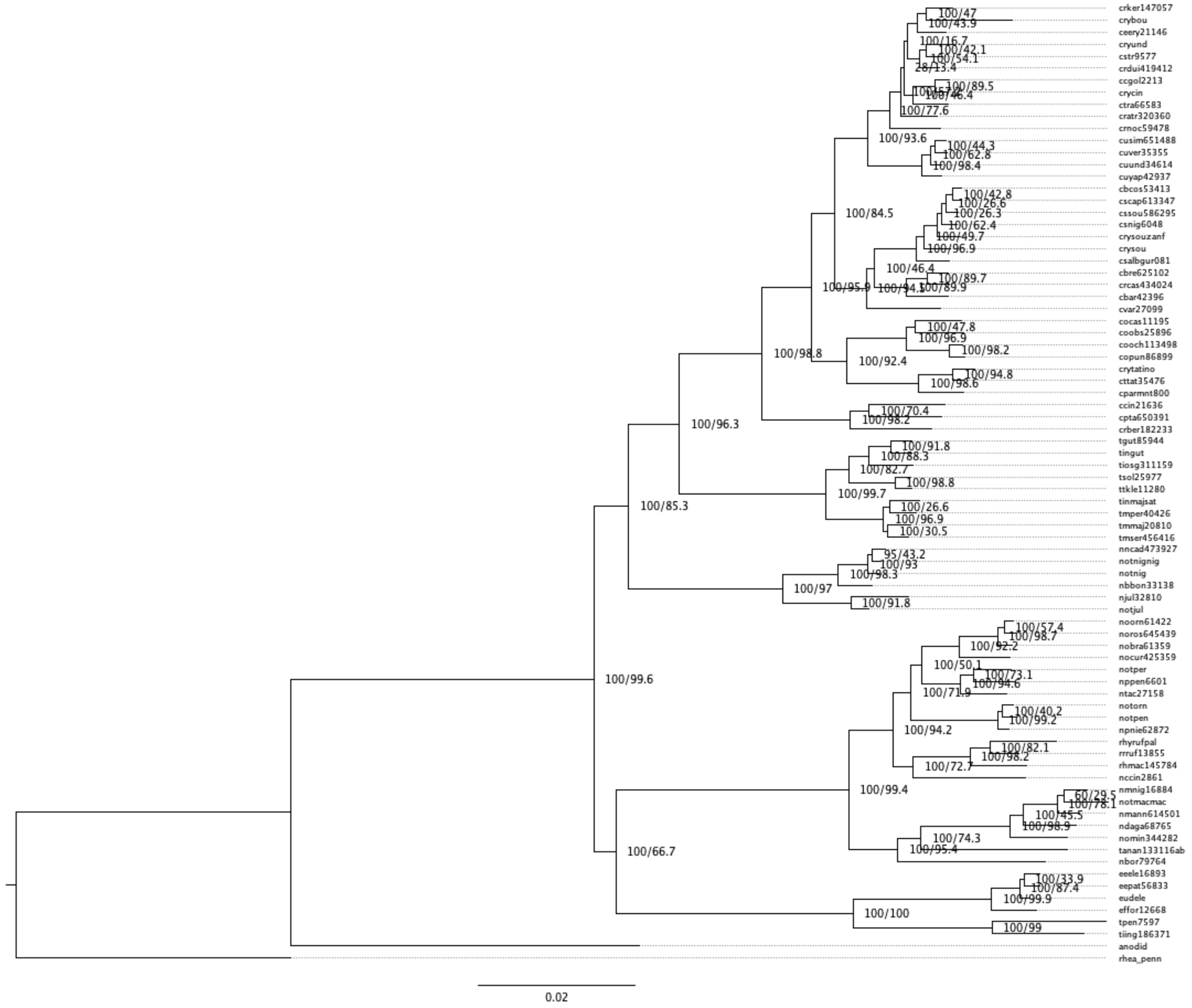

**Figure S19: UCE1000 75% whole dataset iqtree phylogeny**

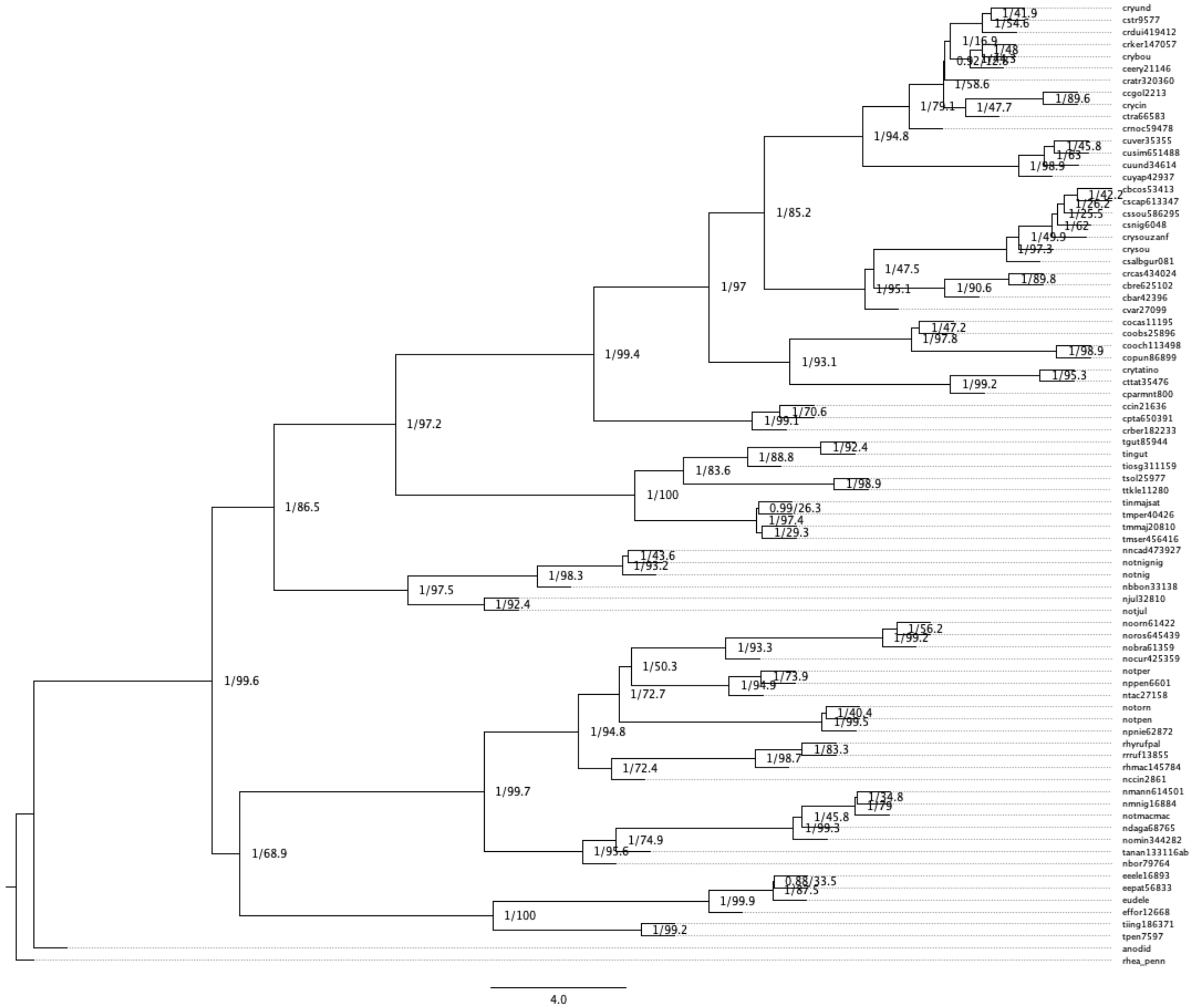

**Figure S20: UCE1000 100% whole dataset Astral phylogeny**

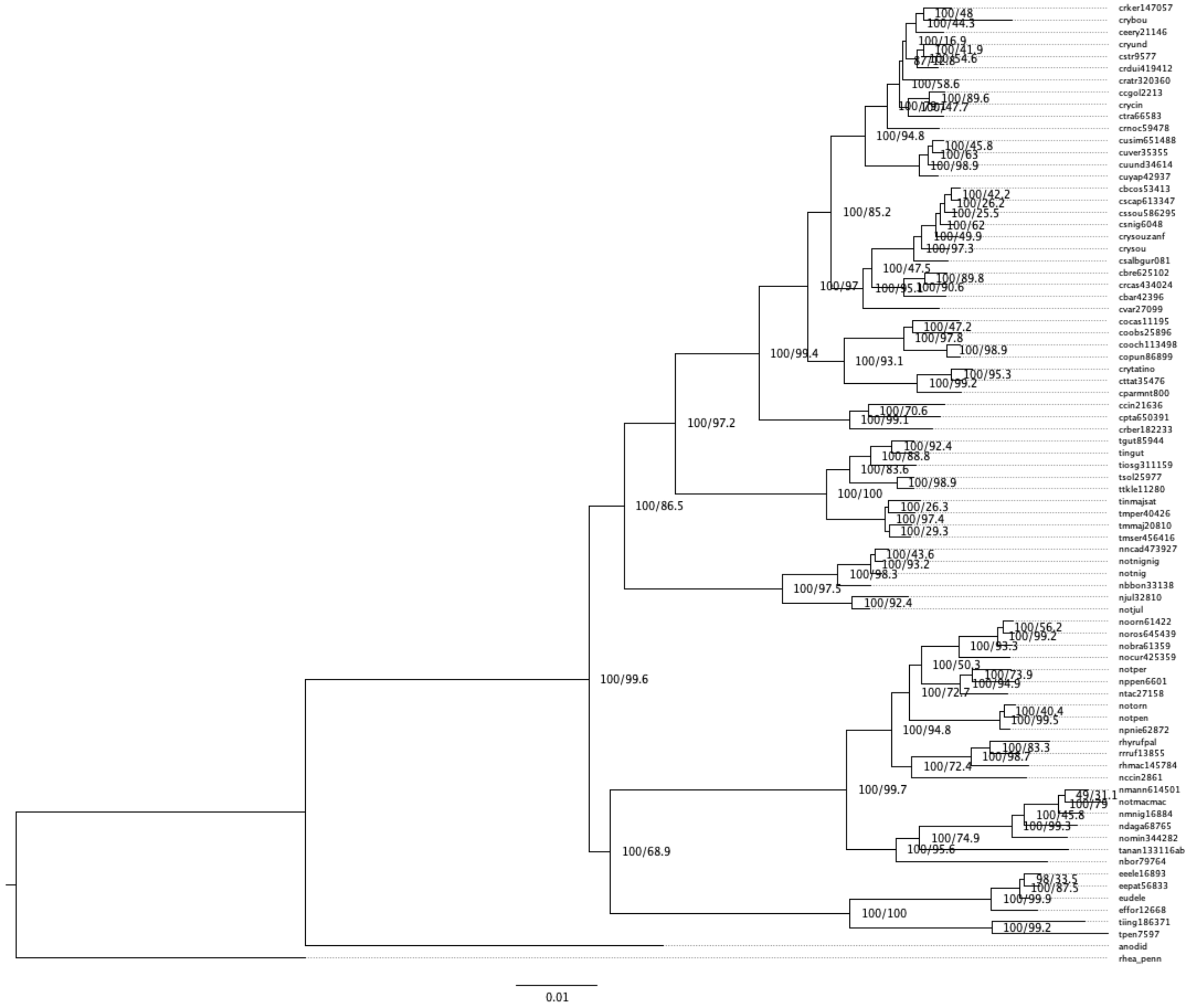


**Figure S21: UCE1000 100% whole dataset iqtree phylogeny**

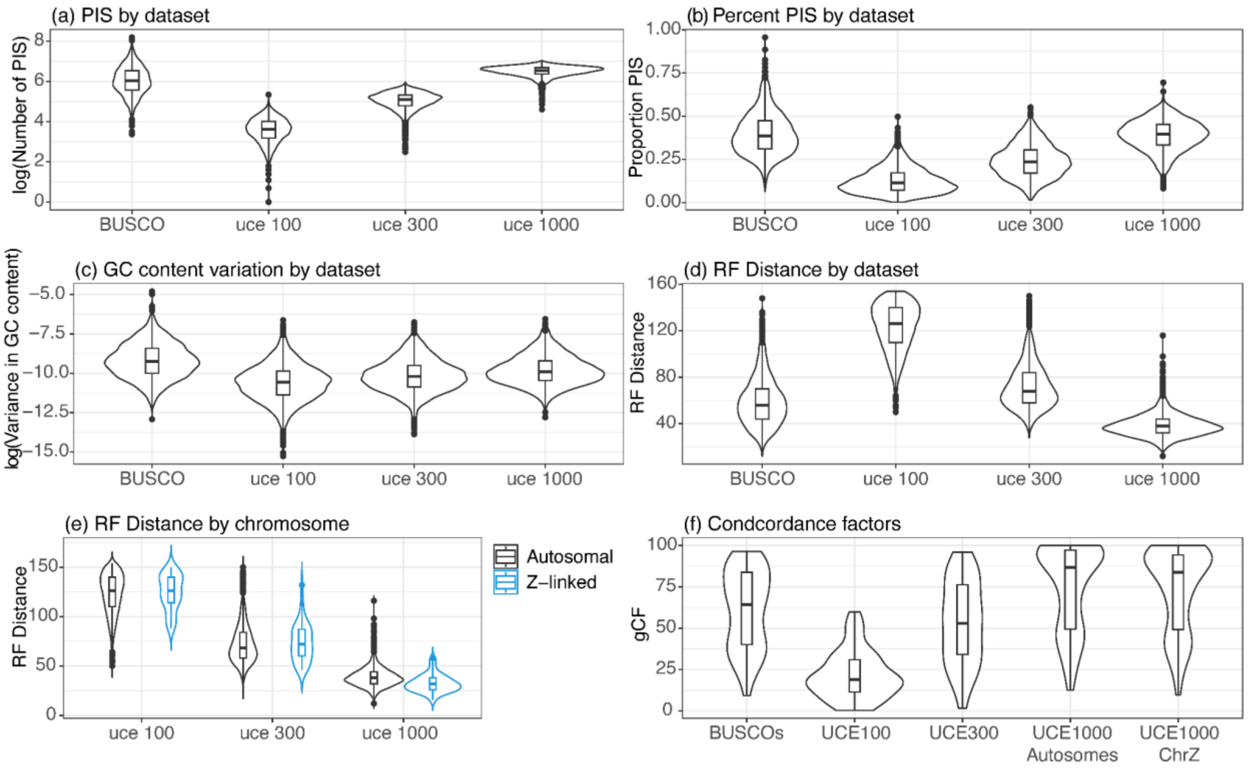


**Figure S22: Information content, variance in GC content, and gene tree heterogeneity for each dataset.** The violin plots show: (a) the ln-normalized number of parsimony informative sites (PIS), (b) the proportion of PIS, (c) the log-normalized variance in GC content, (d) RF distance between gene and species trees, (e) RF distance between gene and species trees with autosomal and Z-linked UCEs shown separately, and (f) concordance factors for all nodes within each MSC species tree.

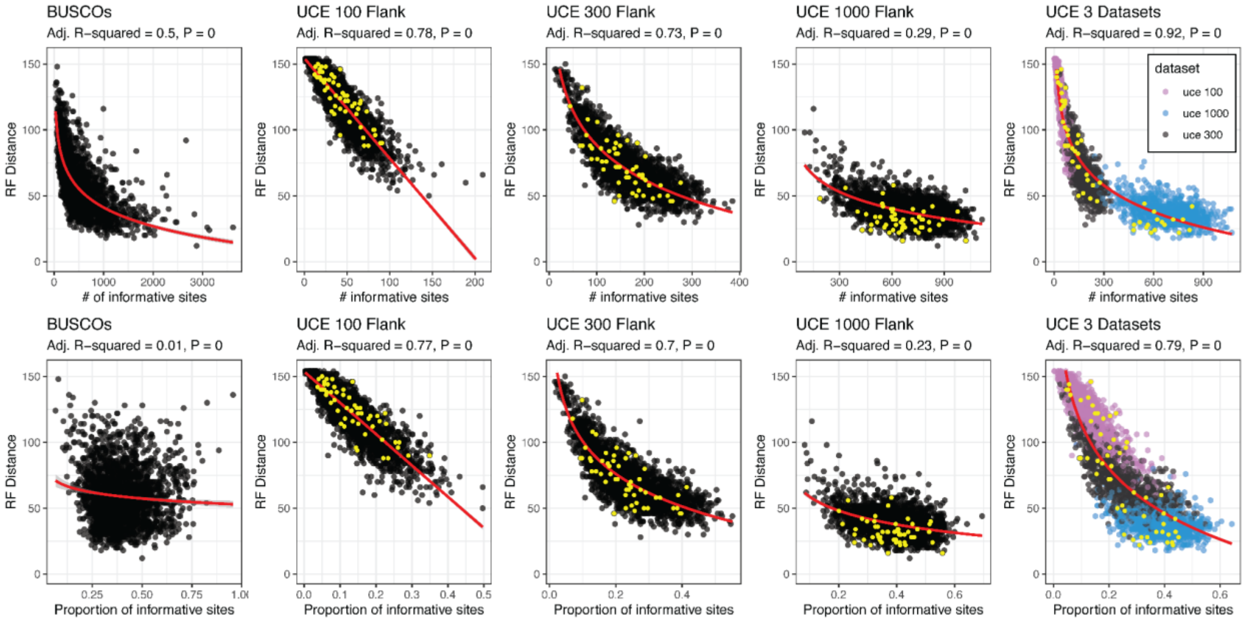


**Figure S23: Covariation between information content and gene tree estimation error.** Linear regression models exploring the relationship between information content and gene tree heterogeneity. The negative correlation in most instances suggests that alignments with less information content result in increased gene tree estimation error. UCEs that map to the Z-chromosome in each dataset are shown in yellow.

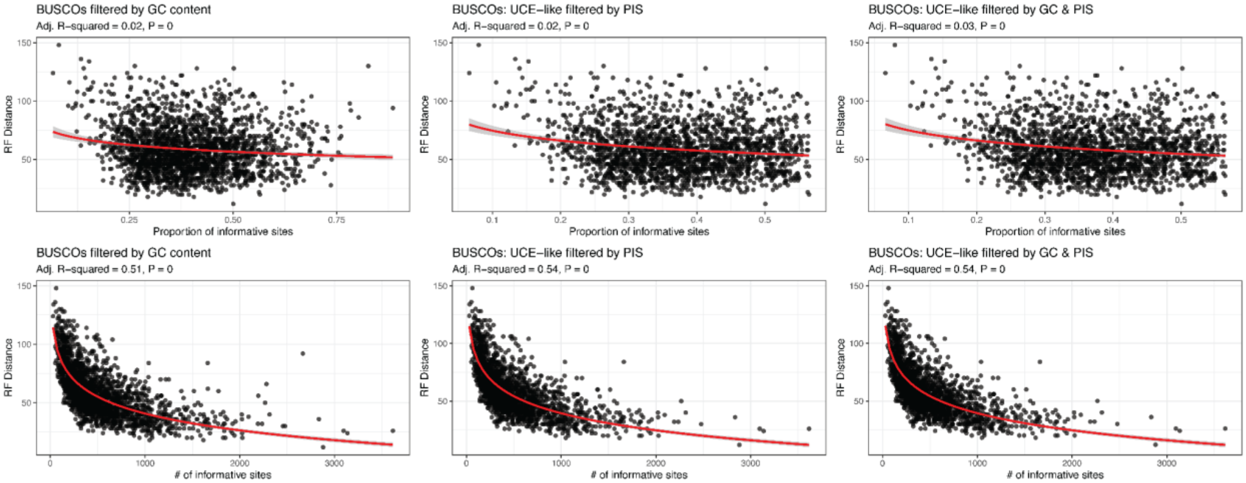


**Figure S24: Linear regressions of information content and RF Distance for filtered BUSCOs.** The BUSCO dataset was filtered three times. First, the extremes of GC content were removed (BUSCOs filtered by GC content). Second, BUSCO's were filtered by the number and proportion of PIS, wherein BUSCOs with greater than two standard deviations away from the mean number and proportion of PIS were removed. Finally, the two BUSCO filters were combined.

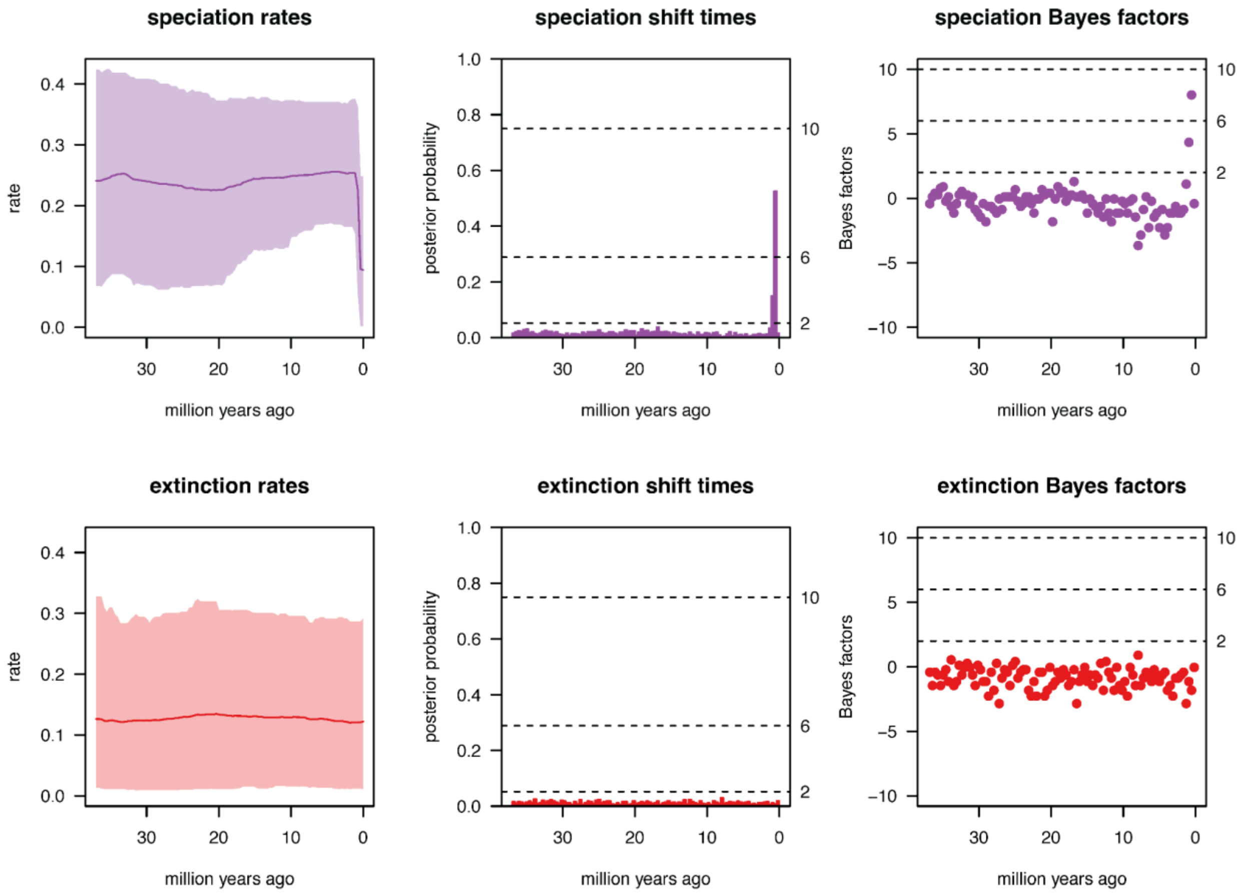


**Figure S25:** Speciation (purple) and extinction (red) times for Tinamidae along with posterior probabilities and Bayes factors for the presence of rate shifts.

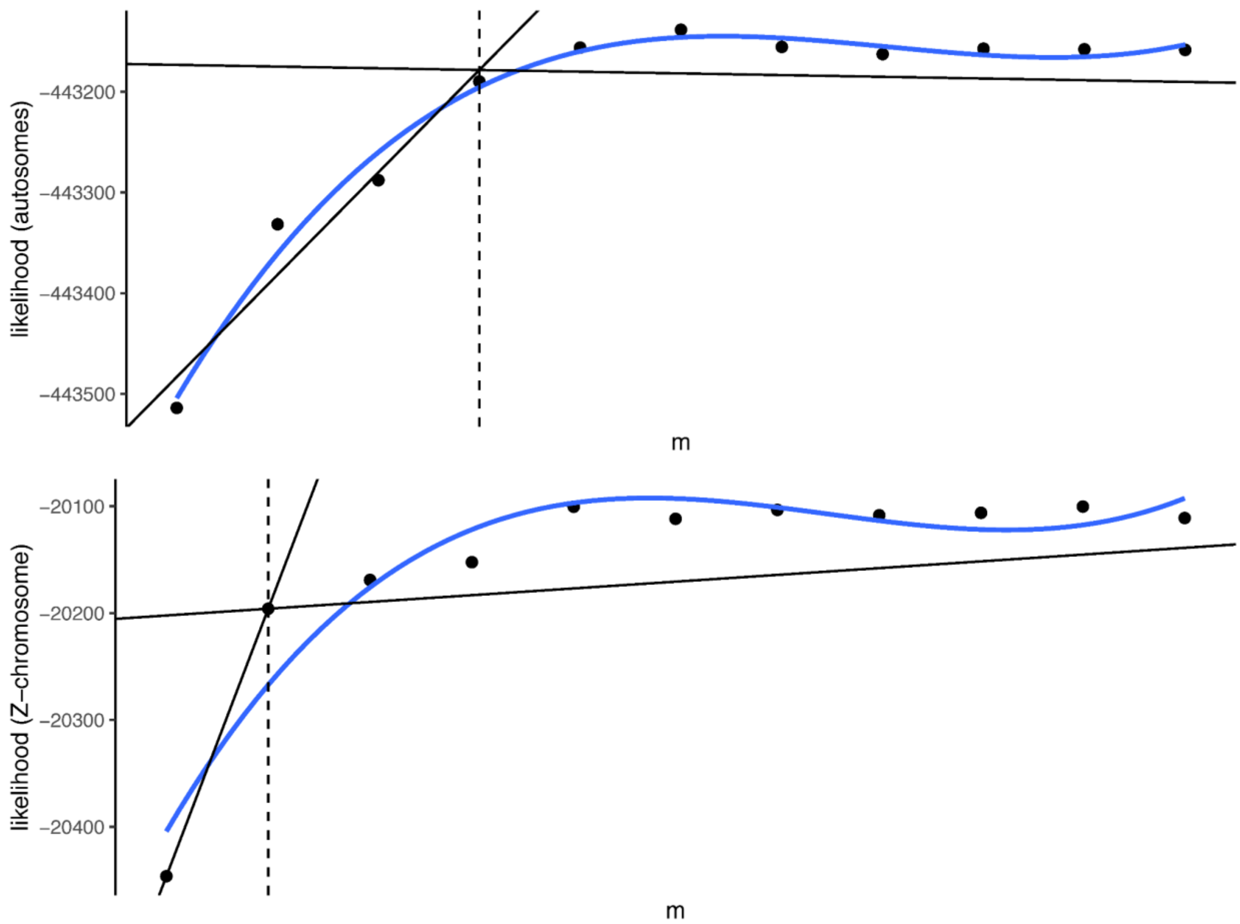


**Figure S26: Breakpoint analysis for the best-fit m-value for the introgression analysis.**

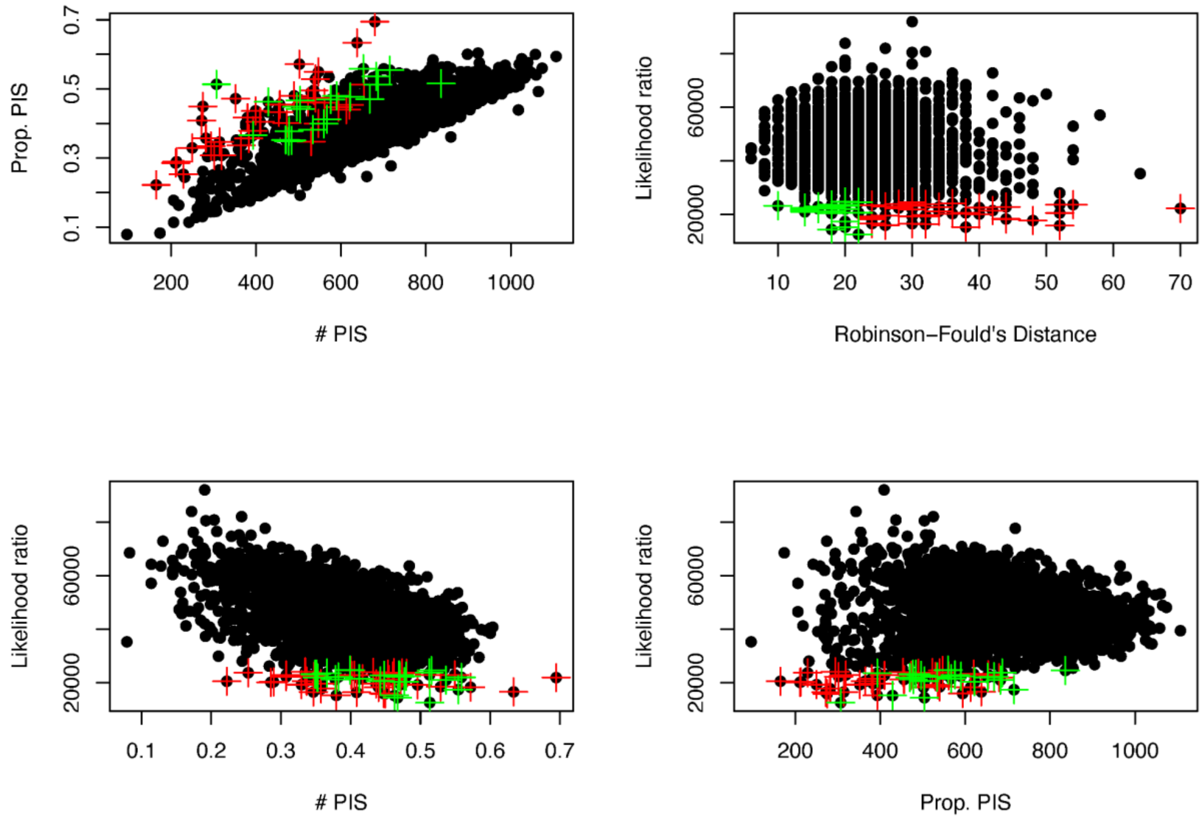


**FigureS27: Results of likelihood ratio and RF filtering for divergence dating analysis.** The plots show values for all autosomal genes before (black points) and after (red and gree crosses) filtering. Red crosses represent the alignment/gene trees two standard deviations below the mean likelihood ratio, and green crosses show those that are also below the mean for RF distance and were retained for use in divergence dating.
